# A haplotype-phased male genome sequence of the stinging nettle, *Urtica dioica* ssp. *dioica*

**DOI:** 10.1101/2025.09.12.675976

**Authors:** Kaede Hirabayashi, Diana Percy, Eric Gonzalez, Michael Deyholos, Quentin Cronk, Marco Todesco

**Author notes:** Corresponding author: Marco Todesco.

## Abstract

Stinging nettle (*Urtica dioica* L.) is a widespread weed of economic significance with a dioecious mating system. Previously, we generated a high-quality genome assembly of a diploid female plant, which showed extreme levels of structural variation between haplotypes. Here, we present a chromosome-level, haplotype-resolved sequence of a diploid male plant; since the male is believed to be the heterogametic sex in *Urtica dioica*, this assembly represents a first step towards elucidating the control of sex determination in this species. This completely independently assembled genome also allows confirmation of previously reported features of the nettle genome, including (1) a high degree of structural variation between haplotypes, including large inversions, (2) the likely existence of polycentric centromeres, and (3) the presence of urticaceous “pain peptide” sequences. Chromosome 8 stands out for its multiple large, nested inversions and high levels of repetitive sequences, features that are often associated with sex determining regions (SDRs). This chromosome is therefore a candidate for further investigations to characterize the sex determination in nettle.

## Background & Summary

The stinging nettle (*Urtica dioica* L. ssp. *dioica*) is a widespread weed common throughout Europe, temperate Asia, and North America. It owes its name to its stinging hairs, which act as a powerful herbivore deterrent. It is often associated with nutrient-enriched human habitats as it has high phosphate demands, ceasing growth when phosphate is limiting, but responding vigorously to its addition^1^. Its natural habitat is probably rich alluvial river floodplains, where its phosphate demands are met by sediment deposition. More recently, *Urtica dioica* has exploited human disturbance and eutrophication to expand as an aggressive ruderal and weedy species. However, stinging nettle has also a long history of positive associations with humans and has been used as a source of medicine^2^, food^3^, and fibre^4^. Ecologically, it acts as a “super-host” for a multitude of invertebrates across Europe^5^. It is a crucial food source for numerous specialist and generalist insects, particularly within the orders Lepidoptera, Coleoptera, and Hemiptera^6^, and for molluscs. Its attractiveness to invertebrate herbivores may be due to its status as a general mineral accumulator, containing high concentrations of calcium, nitrogen, and phosphorus in its tissues^7^.

*Urtica dioica* ssp. *dioica* is a dioecious taxon (i.e. it has separate sexes on different plants). Evidence for male heterogamety was first obtained by Strasburger, who selfed a female plant that had formed some aberrant individual male flowers, obtaining only female progeny^8,9^. Subsequent studies reported complex inheritance patterns and significant maternal effects on seed sex ratios in dioecious *Urtica*, while monoecious individuals produced varying numbers of female flowers based on environmental conditions, suggesting a wholly genetic basis for sex determination in dioecious types but environmental influences in monoecious types^10,11^.

The first published genome sequence of *U. dioica* was the haplotype-resolved assembly of a diploid female individual (clone 11-4)^12^. A haplotype from a tetraploid individual (the more common ploidy level for stinging nettle) was also subsequently generated^13^. Analysis of the diploid female genome assembly revealed several unusual features: (1) Extensive structural variation (SV) between the two haplotypes, particularly on chromosomes 1, 2, 3, and 8; (2) Presence of two types of centromeres, based on the analysis of patterns of repetitive sequences: acrocentric or near telocentric centromeres in five chromosomes (8, 9, 11, 12, 13) and polycentric centromeres in eight others (chr. 1–7, 10). The occurrence of polycentric centromeres had not been previously reported in the Urticaceae family, but is known in the closely related Moraceae; (3) Identification of two copies of a gene putatively encoding the sting peptide urthionin on chromosome 9, suggesting a recent gene duplication. Urthionin is a 42 amino acid-long peptide with cytolytic activity that contributes to nettle’s sting^14^.

Here, we report a chromosome-level, haplotype-resolved genome assembly for a diploid male individual, the putative heterogametic sex of stinging nettle. In this assembly, we observe many of the same genomic features mentioned above for the female stinging nettle genome assembly^12^, confirming them and validating the correctness and accuracy of this new male stinging nettle genome assembly. This male assembly will be foundational to further studies aimed at characterizing the sex-determining region (SDR) in *Urtica* and determining the cause of the male heterogamety. We provide, for each chromosome, information on which haplotype is maternally- or paternally-derived, taking advantage of the fact that the male individual that we sequenced is the offspring of the female individual for which a genome assembly has been recently released^12^. This knowledge will help future efforts aimed at identifying the SDR of stinging nettle. In particular, a highly variable region in chromosome 8 presents several of the hallmarks of an SDR, including complex large inversions and a possible 8 Mbp insertion in the paternally-inherited haplotype, and will be a promising candidate for further studies on sex determination in stinging nettle.

## Methods

### Plant materials and sequencing

We selected a male plant (U48) from a progeny panel derived from a cross between two wild-collected *U. dioica* diploid cytotypes: 11-4 (female, River Jiu, north of Rovinari, Romania, for which we previously produced a genome assembly)^12^ and 7-5 (male, River Struma, north of Boboshevo, Bulgaria)^15^. A voucher specimen has been deposited in the herbarium of the Beaty Biodiversity Museum at the University of British Columbia (UBC). Leaf tissue was frozen in liquid nitrogen and high-molecular weight DNA was isolated using a modified CTAB method^16^. A PacBio HiFi library was constructed and sequenced on a PacBio Revio instrument at Canada’s Michael Smith Genome Sciences Centre (GSC) in Vancouver, BC, Canada, generating a total of 3.23 million HiFi reads (∼77.5 Gbp; ∼129X genome-wide coverage) with a mean quality of Q32 and mean length of 24,025 bp. Quality control of HiFi reads was performed using SMRT Link v13.1 with the command runqc-reports v10.4.6 (https://www.pacb.com/support/software-downloads/). We also prepared a Hi-C library according to the method described in ^12^. The Hi-C library was sequenced with PE150 reads on an Illumina NovaSeq X Plus instrument at GSC, producing 108 million paired Hi-C reads (32.2 Gbp; ∼54X genome-wide coverage).

#### *De novo* genome assembly and quality assessment

Raw HiFi reads were filtered for >Q20 using fastq-filter v0.3.0^17^, resulting in 88.2% of the HiFi reads (∼113X genome coverage) being retained and used for the *de novo* assembly. A preliminary k-mer analysis was performed on these reads using Genomescope v1.0^18^ to determine expected heterozygosity. Hi-C reads were quality-checked using fastqc v0.12.1^19^ and were not filtered or trimmed at this stage. To produce the draft haplotype-resolved genome assembly, we used Hifiasm v0.19.8-r603^20^ with HiFi and Hi-C reads as input files. Tuning parameters such as --hom-cov and -s did not improve the quality of the initial assembly; therefore, we proceeded with the assembly produced using the default settings.

We used Hi-C data to scaffold the two haplotype assemblies (male haplotype 1: MH1; male haplotype 2: MH2) independently, following the YaHS pipeline v1.2.2^21^, which combines assisted scaffolding and manual examination of the contigs. In brief, Hi-C reads were mapped to the draft genome assembly using Juicer v1.9.9^22^. The resulting merged_nodups.txt file was then converted to a bed file using a custom awk script where each line has 1) contig ID a read was mapped to, 2) start of the alignment, 3) end of the alignment, 4) read ID with /1 and /2 denoting a paired read, and 5) mapping quality^23^. To avoid inaccurate alignment for the initial scaffolding process, we retained and visualized only read alignments with mapping quality above 0. Next, the draft genome was scaffolded with YaHS’s default parameters. We then manually examined the scaffolds for possible misassemblies using Juicebox v1.11.08^24^, according to the YaHS manual curation pipeline, and assigned chromosomal boundaries. To further correct any switch errors (i.e. contigs that were placed in the wrong haplotype), we mapped Hi-C reads to the combined MH1+MH2 assembly using Juicer and converted the merged_nodups.txt to Juicebox compatible files using the 3D-DNA pipeline v189022^25^ with no mapping quality filter (parameter -q 0)^25^. Although allowing mapping quality 0 reads is generally not recommended because it can lead to erroneous alignment, we found that this significantly improved our ability to resolve repetitive regions when aligning Hi-C reads to both haplotypes at the same time. While we identified no obvious switch errors, we observed many small unassigned contigs (∼100-200 kbp) that showed a strong association with a particular chromosome’s centromeric or telomeric region. We positioned these contigs within the chromosome scaffolds manually, provided that the linear interaction pattern within the chromosome remained continuous after their insertion. Additionally, a large contig, which appeared to be mainly composed of a repetitive region, showed strong interactions only with the centre of chromosome 8 of MH2, and was manually inserted into the existing gap between the contigs within that chromosome. While these manual curation steps caused a noticeable decrease in contig N50 and scaffold N50, they produced a cleaner Hi-C interaction heatmap and haplotype assemblies that are likely to be more complete (Supplementary Fig. 1).

Lastly, we used purge_dups v1.2.5^26^ to remove duplicates and small contigs/unassembled reads that could not be placed within the main scaffolds. For this step, manual cutoff values (MH1: 5 48 78 96 158 360; MH2: 5 59 59 60 155 360) were estimated based on the k-mer histogram produced from the >Q20 PacBio HiFi data, according to the purge_dups guidelines. An additional round of manual curation was performed using Juicer, YaHS, and Juicebox to make sure that both haplotype assemblies did not contain any obvious misassembly. Chromosome numbers and orientations in the finalized assemblies were then modified to match those of the female haplotype-resolved assembly^12^. Assembly quality was assessed using BBMap v39.06^27^ and BUSCO scores (dataset: eudicot_odb10) with BUSCO v5.1.2^28,29^.

The final scaffolded assemblies for MH1 and MH2 had a total length of 576,923,711 bp and 536,126,483 bp, consisting of 833 and 79 scaffolds (N50 = 45.3 Mbp and 49.8 Mbp), respectively; 90.1% and 99.2% of the total assembly length were scaffolded inside the 13 chromosomes of the stinging nettle genome. These assembly sizes are consistent with previous estimates of genome size for stinging nettle^15^, and with the size of the published female stinging nettle assembly^12^. BUSCO scores for the final assemblies showed high completeness (C ≈ 93%) and low duplication rates (D ≤3%; Table 1; Supplementary Table 1). Despite its relatively small size, the genome of stinging nettle contains a high amount of structural variants, especially on chromosomes 1, 2, 3, 6, and 8 (Fig. 1a); with the exception of chromosome 8, these structural variants coincide with regions of low gene density (Fig. 1a,b). To compare the two haplotypes at the nucleotide level, we used SyRI v1.7.0^30^ and visualized the output synteny and structural variations, including inversions, duplications, and translocations, using Plotsr^31^ (Supplementary Fig. 2).

**Table 1:**
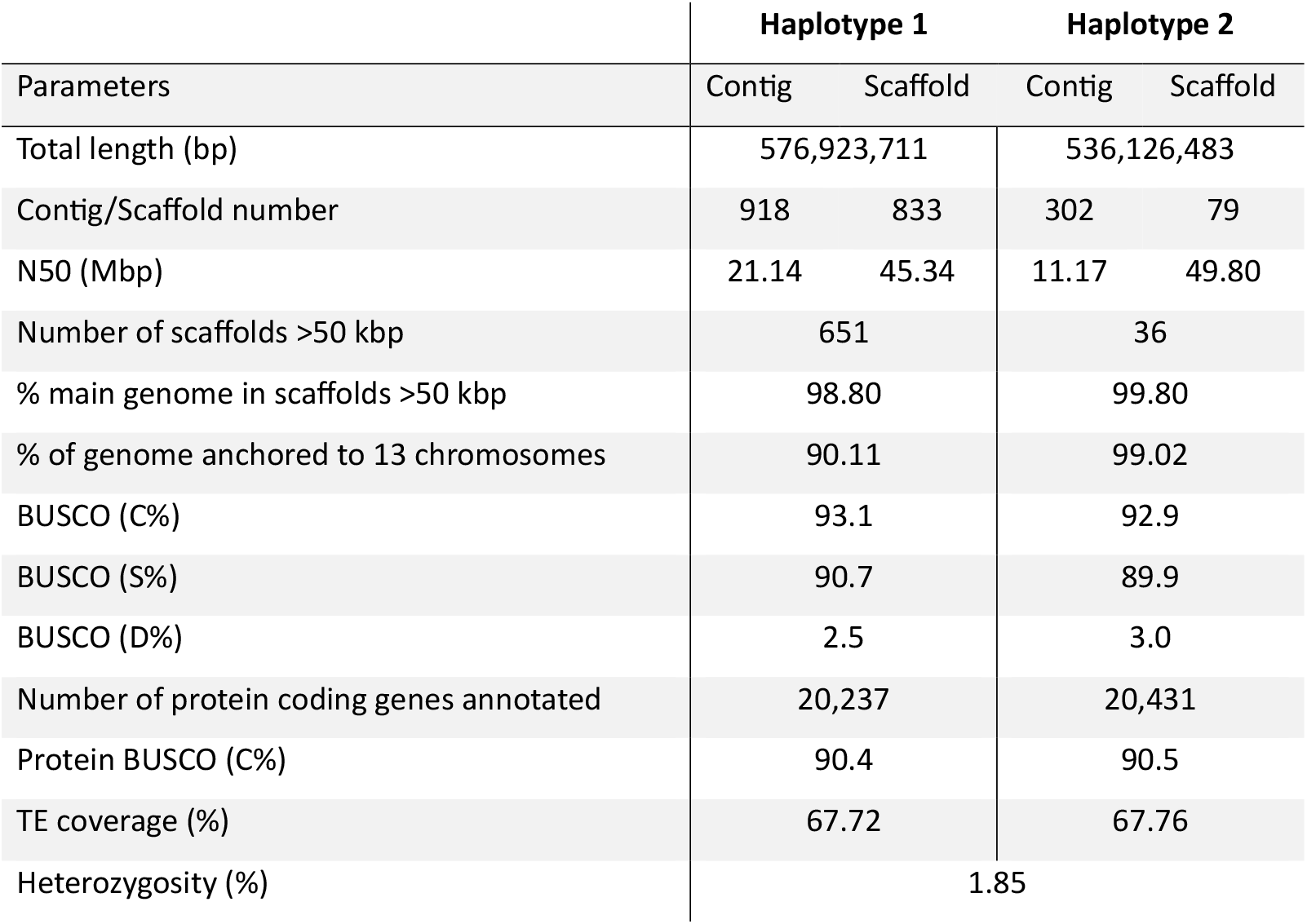
Genome assembly statistics.

|  | Haplotype 1 |  | Haplotype 2 |  |
| --- | --- | --- | --- | --- |
| Parameters | Contig | Scaffold | Contig | Scaffold |
| Total length (bp) | 576,923,711 |  | 536,126,483 |  |
| Contig/Scaffold number | 918 | 833 | 302 | 79 |
| N50 (Mbp) | 21.14 | 45.34 | 11.17 | 49.80 |
| Number of scaffolds >50 kbp | 651 |  | 36 |  |
| % main genome in scaffolds >50 kbp | 98.80 |  | 99.80 |  |
| % of genome anchored to 13 chromosomes | 90.11 |  | 99.02 |  |
| BUSCO (C%) | 93.1 |  | 92.9 |  |
| BUSCO (S%) | 90.7 |  | 89.9 |  |
| BUSCO (D%) | 2.5 |  | 3.0 |  |
| Number of protein coding genes annotated | 20,237 |  | 20,431 |  |
| Protein BUSCO (C%) | 90.4 |  | 90.5 |  |
| TE coverage (%) | 67.72 |  | 67.76 |  |
| Heterozygosity (%) | 1.85 |  |  |  |

**Figure 1:**
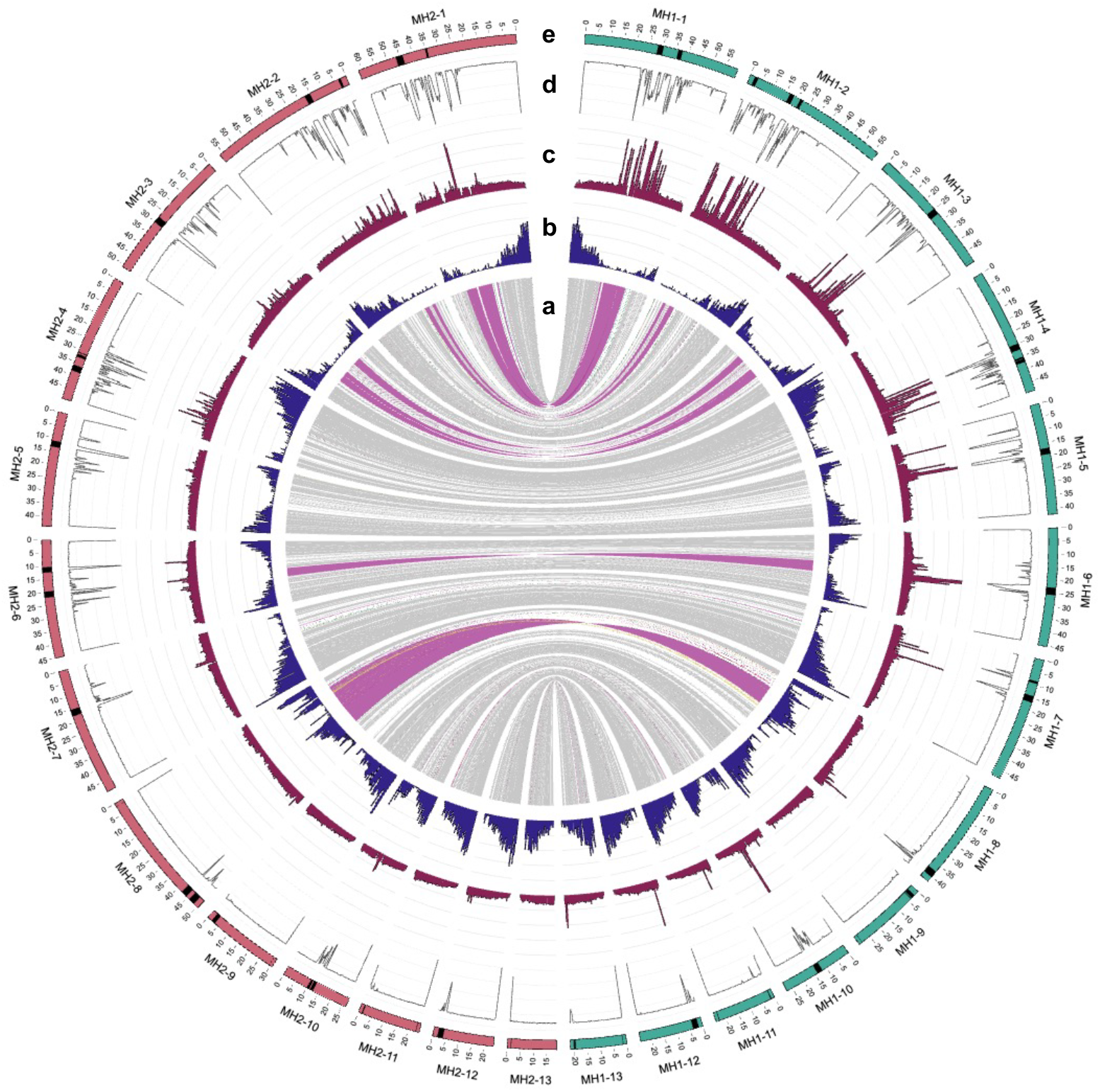
Circos plot for the male nettle genome assembly. From the inside out, tracks show a) SVs identified by SyRI; grey = syntenic regions, purple = inversions, green = translocations, blue = duplications, b) gene density in 500 kbp windows, c) TEs density in 500 kbp windows, d) Shannon diversity of repeats mean score per 250 kbp window, e) putative centromeric locations in black bands based on RepeatOBserver Shannon diversity score. Haplotype 1 is in green on the right, haplotype 2 is pink on the left. The maximum y-axis of the TE counts was set to 4000 to show variation across the genome, resulting in a total of 25 windows that exceed the plot’s range. TE counts for these windows with >4000 TEs are reported in Supplementary Table 4.

### Gene and repeat annotations

We performed whole genome gene and repeat annotation using BRAKER3 v3.0.8^32,33^ and Extensive Denovo TE Annotator v2.2.1^34^ pipelines, respectively. To do so, we first soft-masked the genome with the Red command: redmask.py v0.0.2^35^. Publicly available RNAseq reads from *Urtica dioica*^36^ were downloaded and filtered with Trimmomatic v0.39 (parameters: ILLUMINACLIP: TruSeq3-PE.fa:2:30:10:2:True SLIDINGWINDOW:4:15 LEADING:3 TRAILING:3 MINLEN:36)^37^, then aligned to our genome using Hisat2 v2.2.1^38^. The resulting output was then converted to BAM format using samtools v1.17^39^. Using the AUGUSTUS parameters that we previously generated for the female nettle assembly resulted in a less complete annotation of the male assembly. Therefore, we ran the braker.pl command with the new species flag -- species Urtica_dioica_male to train AUGUSTUS, with --prot_seq ‘Viridiplantae’ dataset downloaded from OrthoDB^40,41^, --bam RNAseq alignment file, and --softmasking options. This means that the male MH1 was annotated *de novo* and independently of the female assembly annotation data. To annotate the male MH2 haplotype, we used the same --species Urtica_dioica_male with --useexisting flag to carry over the AUGUSTUS training parameters from MH1. For TE annotation, we ran EDTA with default parameters. Annotation of putative pain peptides was performed as previously described^12^, based on the peptide sequences published in ^14,42^. The results of orthologous hits are presented in Supplementary Table 2.

Gene annotation identified 20,237 unique genes and 22,384 transcripts in MH1, with protein BUSCO scores of 90.4 % (C), 79.7 % (S), and 10.7 % (D). Similarly, MH2 annotation contained 20,431 unique genes and 22,607 transcripts, with BUSCO scores of 90.5 % (C), 79.4 % (S), 11.2 % (D) (Table 1, Supplementary Table 3). We identified 724,918 and 572,024 TEs spanning 390,681,099 bp (67.72 %) and 363,225,467 bp (67.76 %) of the genome in MH1 and MH2, respectively. The most abundant type of TEs was Long Terminal Repeat (LTR) retrotransposons (MH1: 47.67 % and MH2: 46.04 %), followed by Terminal Inverted Repeats (TIR; MH1: 8.70 %, MH2: 15.08 %) (Supplementary Table 4).

We used RepeatObserver v0.1.0^43^ with default cut-off values to identify putative centromeric regions and to generate Fourier heatmaps for each chromosome. Analysis of tandem repeats using RepeatOBserver is consistent with the presence of polycentric chromosomes in the nettle genome (chromosomes 1-7), alongside metacentric (chromosome 10) and acrocentric or near telomeric centromeres (chromosomes 8, 9, 11, 12, 13; Supplementary Fig. 3, Supplementary Table 5). A circos plot^44^ was generated to visualize gene/repeat counts per 500 kbp windows, as well as the mean Shannon diversity scores from RepeatOBserver averaged per 250 kbp windows (Fig. 1).

### Parental assignment of haplotypes

We took advantage of the fact that the female individual sequenced in ^12^ is the mother of the newly sequenced male individual to identify which of its haplotypes are maternally- or paternally-derived. We used minimap2 v2.24-r1122^45^ to perform pairwise alignments of four haplotypes from the *U. dioica* ssp. *dioica* female^12^ and male (this study) individuals, denoted as female H1 (FH1), female H2 (FH2), male H1 (MH1), male H2 (MH2), to the two female haplotypes (FH1 and FH2)^44^. We expected one of the haplotypes of the male assembly to be identical to one of the haplotypes of the female assembly, or to be a mix of the two female haplotypes in cases where recombination has occurred between them. To verify this, we filtered the alignment file between female and male haplotypes to retain hits with quality >30 and calculated the percent identity of the nucleotide match as (number of matching nucleotides / alignment length) x 100 for each alignment detected. We then plotted the alignment results filtered by ≥95% matching nucleotides to identify the maternally-inherited (and, by exclusion, the paternally-inherited) chromosomes in the male nettle assembly (Fig. 2, Supplementary Fig. 4). In cases where there appear to be no recombination (chromosomes 1 – 3, 6 – 8), near-perfect alignment was observed between one of the female haplotypes and one of the male haplotypes, but not in pairwise comparisons with the other male haplotype. For example, in chromosome 8 (Fig. 2), only MH1 has a complete, identical nucleotide match to FH1, while MH2 does not align perfectly to FH2. The MH2 haplotype does not align perfectly to either FH1 or FH2. This indicates that, for chromosome 8, MH1 is maternally-derived and MH2 is paternally-derived. This analysis also identified multiple recombination events, as shown for instance in chromosome 12 (Fig. 2). We observed a clear indication of one recombination event each near the extremities of chromosomes 4, 5, 10, 11, and 12, where high similarity of the male haplotype switches from one to the other female haplotype (Supplementary Fig. 4). A larger recombined region, corresponding to more than one-third of the chromosome, was found in chromosome 9. While most recombination events were easily detectable, a few possible cases were difficult to assign confidently based only on the alignment percentage distribution. For example, a recombination seemed to have occurred at the beginning of chromosome 13, but in the putatively recombined region percent identity is high with either FH1 or FH2 (∼97%). We also note that a percent identity was slightly lower around centromeric regions, possibly due to poor alignment caused by the presence of extensive repeats. Contrary to expectations^46^ we did not observe recombination in several chromosomes. Intriguingly, recombination events were rarer in chromosomes with predicted polycentric centromeres (two recombination events in seven chromosomes) than in chromosomes with metacentric or acrocentric ones (five recombination events in six chromosomes). While based on very limited data (one generation of recombination in a single individual), this observation is consistent with known patterns of reduced recombination in holocentric chromosomes^47^. The inferred maternal and paternal chromosome assignment is summarised in Supplementary Table 6. To quantify how much of each chromosome was inherited from the corresponding maternal haplotype in the male offspring, we filtered the alignment file to retain only alignments with nucleotide match >99.7% and divided the sum of the length of these alignments in each chromosome by the total length of that chromosome (= total alignment length per chromosome / chromosome length x 100; Supplementary Table 6).

**Figure 2:**
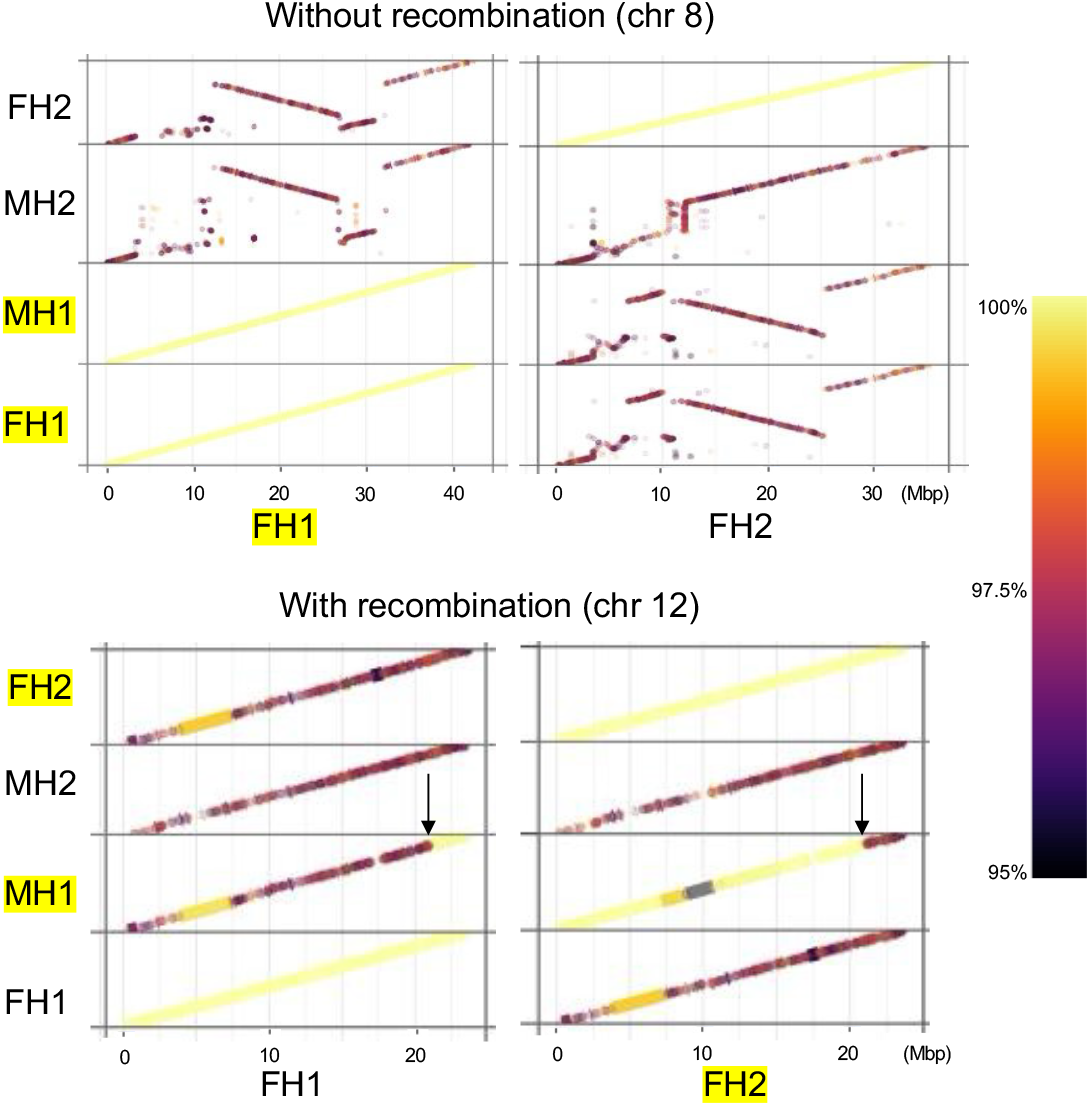
Comparison of four haplotypes; FH1 = female haplotype 1, FH2 = female haplotype 2, MH1 = male haplotype 1, MH2 = male haplotype 2. Pairwise alignments at the nucleotide level using FH1 (left) and FH2 (right) as a reference. Matching chromosome pairs, are highlighted in yellow. An example of parental assignment in the absence (top) and presence of visible recombination events (bottom) is shown. The alignments are coloured by nucleotide match in percent identity, and only the alignment with an identity of more than 95% is shown as a line. The lighter the colour, the higher the percent identity (i.e., yellow = 100%, dark purple = 95%). Pair of male and female haplotypes that have the highest similarity, representing the maternally-inherited male haplotype and its corresponding haplotype in the female assembly, are highlighted in yellow. The full list of chromosome assignments is found in Supplementary Table 6, and plots for all chromosomes are found in Supplementary Fig. 4.

## Data records

The raw sequence data (PacBio HiFi, Illumina Hi-C) reported in this paper have been deposited in the Sequence Read Archive (SRA) of the National Center for Biotechnology Information (NCBI, https://www.ncbi.nlm.nih.gov) under the accession numbers SRR35053641 (PacBio HiFi) and SRR35053640 (Hi-C) with the BioProject accession PRJNA1308592, BioSample accession SAMN50702769. The assembled genomes have been deposited in the NCBI GenBank under the GenBank accessions JBQRAE000000000 (MH1; PRJNA1308592), JBQRAF000000000 (MH2; PRJNA1308630). We have also deposited the corresponding files, including genome assemblies, genome annotation, and TE annotation files, at Figshare: https://figshare.com/projects/A_high-quality_phased_genome_assembly_of_stinging_nettle_Urtica_dioica_ssp_dioica/230981.

## Technical validation

The availability of a previous, independently generated diploid assembly for a female stinging nettle individual allows direct validations of many of the features of the male assembly presented here. As mentioned above, we performed pairwise comparisons between the two male and the two female haplotypes (Fig. 2; Supplementary Fig. 4). Additionally, we used GENESPACE v1.3.1^48^ to independently compare synteny between these haplotypes based on gene order, by using the transcripts annotated on the four haplotypes as input files (Fig. 3a). To run GENESPACE, we used OrthoFinder v2.5.4^49^ to find orthogroups and then used MCScanX^50^ to find synteny blocks. These analyses confirmed that most of the regions of high structural variability found between haplotypes in the male assembly correspond to regions of high structural variability in comparisons with and between the female haplotypes presented in ^12^. This supports the correctness of the male assembly and further highlights the large extent of interspecific structural variation found in stinging nettle. It also identifies a highly variable region of chromosome 8 as a candidate for the SDR in stinging nettle; this region displays multiple multi-Mbp chromosomal inversions and other rearrangements, including a possible 8 Mbp insertion in the paternally-inherited haplotype that is not observed in the maternal haplotypes (Fig. 3b). Additionally, comparison of RepeatOBserver analyses found that the same putative centromere type (polycentric, metacentric, acrocentric) was predicted for the different chromosomes between male and female assemblies of stinging nettle. Finally, two copies of genes putatively encoding the pain peptide urthionin A^14^ were found on chromosome 9 in both haplotypes (Supplementary Table 2), as is the case in the female stinging nettle genome assembly.

**Figure 3:**
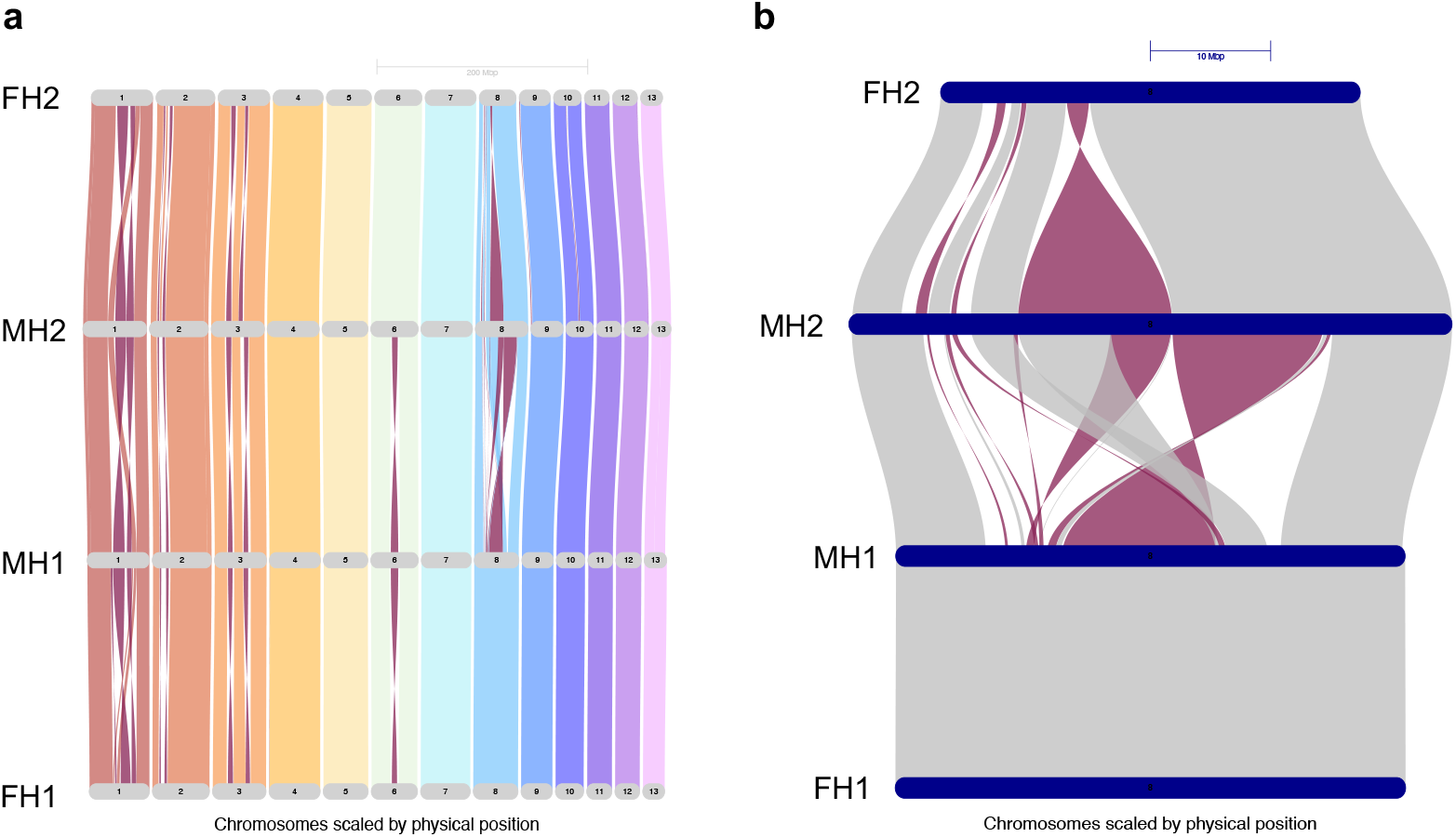
Riparian plot for the four haplotypes showing a) all 13 chromosomes and b) chromosome 8. The plot was created in GENESPACE by comparing the order of gene blocks across the haplotypes (blkSize = 5, blkRadius = 5). Purple = inverted blocks.

## Data Availability

Raw sequencing data and assembled genomes are available at NCBI under BioProject/accession numbers PRJNA1308592/JBQRAE000000000 (haplotype 1; MH1) and PRJNA1308630/JBQRAF000000000 (haplotype 2; MH2). The male plant used in this study has the BioSample ID: SAMN50702769, and is vouchered as a specimen in the herbarium at UBC Beaty Biodiversity Museum. PacBio HiFi reads and Illumina Hi-C reads are provided in .fastq format at the Sequence Read Archive with accession numbers SRR35053641 and SRR35053640, respectively. The genome assemblies and the genome annotations are also deposited in Figshare: https://figshare.com/projects/A_high-quality_phased_genome_assembly_of_stinging_nettle_Urtica_dioica_ssp_dioica/230981.

## Code availability

All the code used for this project is an extension from previous work^12^ and is found at https://github.com/kaede0e/stinging_nettle_genome_assembly. This includes in particular: the full pipeline for the assembly presented in this paper in 1_genome_assembly_with_PBHiFi_HiC/how_to_run_assembly_workflow_v2.md; the gene annotation pipeline in 2_annotation/how_to_run_BRAKER3_with_conda.sh; the R script for pairwise haplotype alignment visualization in 3_comparative_genomics/PLOT_cont2_allCHR_PER.R; and the coupled calculation for maternal chromosome assignment in 3_comparative_genomics/calculate_chromosome_percent_inheritance.sh.

## Author contributions

Conceptualization: QC and MD; funding and resources: QC, DP and MT; data production: KH; formal analyses, investigation, and visualization, KH, EG, and MT; sample collection: DP and QC; genotype maintenance and crossing, QC; sample preparation and laboratory work: KH; writing, review and editing: all authors. All the authors read and approved the final manuscript.

## Acknowledgments

We thank the Digital Research Alliance of Canada (DRAC) for access to their computational resources. We acknowledge the following funding sources: Natural Science and Engineering Research Council (NSERC) Discovery Grants to MT (RGPIN-2023-03344) and QC (RGPIN-2019-04041), and the Natural History Museum, London (UK) for funding to DP for fieldwork in Europe.

## Supplementary information

Supplementary Figures 1-4

Supplementary Tables 1-6

