## Supplementary Fig. for "A haplotype-phased male genome sequence of the stinging nettle, *Urtica dioica* ssp. *dioica*"

**Supplementary Figures**

**Supplementary Fig. 1: Hi-C contact map for the male stinging nettle genome assembly.**

**Supplementary Fig. 2: Structural comparison between MH1 and MH2.**

**Supplementary Fig. 3: Genome-wide patterns of tandem repeats.**

**Supplementary Fig. 4: Maternal and paternal chromosome assignment.**

Urtica genome

a)

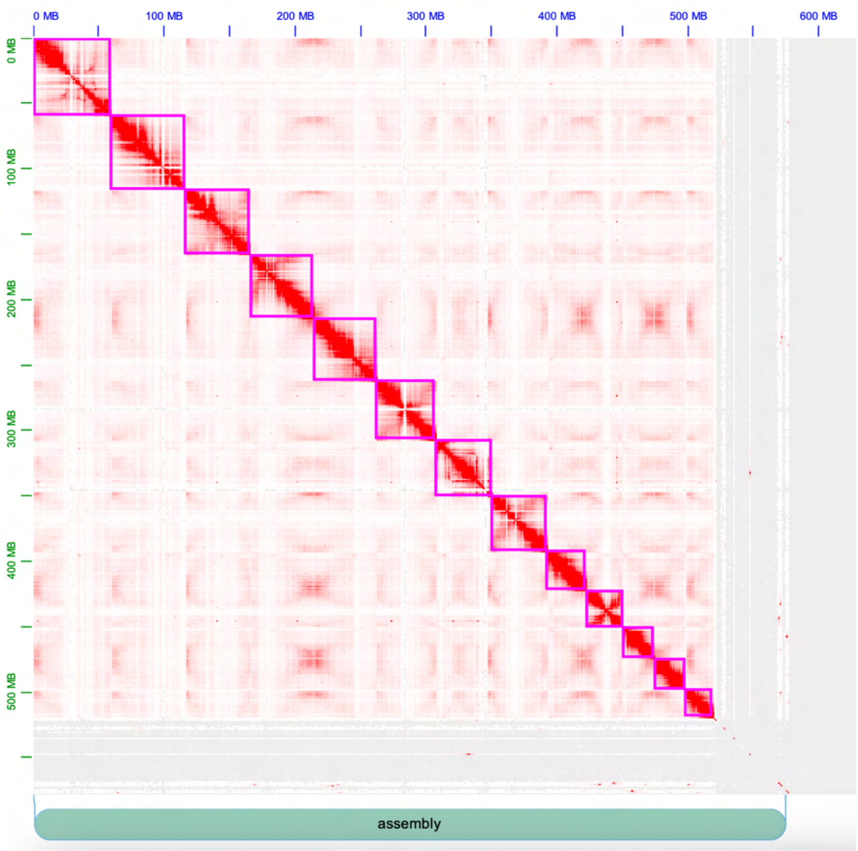


b)

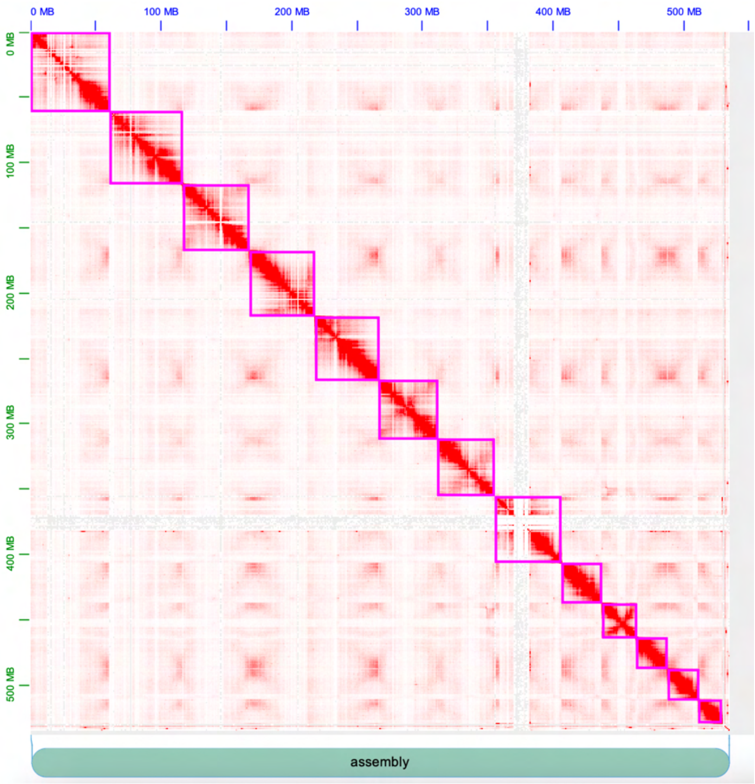

c)

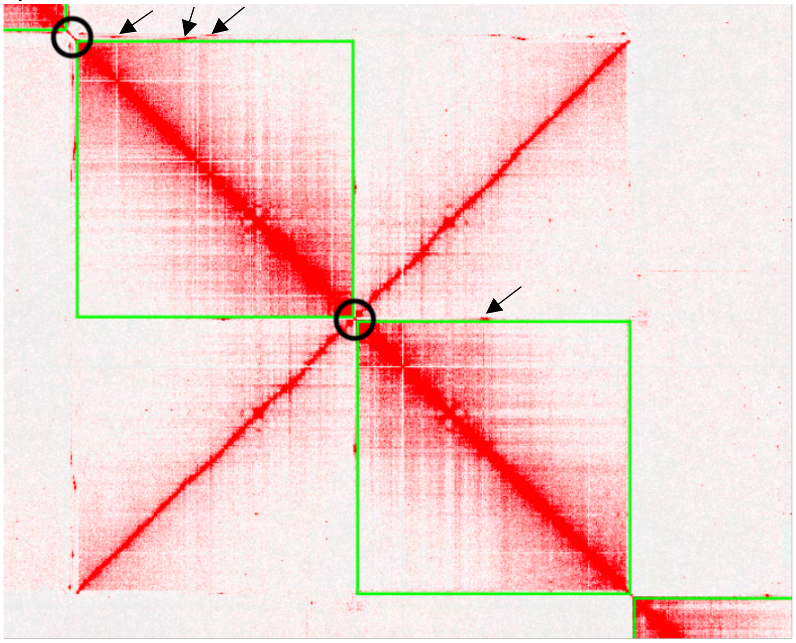

**Supplementary Fig. 1: Hi-C contact map for the male stinging nettle genome assembly.** Hi-C heatmap of the final assemblies for a) haplotype 1 – MH1, and b) haplotype 2 – MH2, of the *Urtica dioica* ssp. *dioica* male diploid genome. In (a) and (b), purple lines are manually assigned chromosome boundaries. c) An example of un-scaffolded contigs (circled in black) that we inserted in the corresponding chromosomes at telomeric sites and centromeric sites based on significant interactions (black arrows). In (c), Hi-C reads are mapped against MH1 and MH2 simultaneously, and all mapped reads are shown (mapping quality ≥ 0 are shown). MH1 (upper left quadrant) and MH2 (lower right quadrant) are plotted in opposite orientations for chromosome 12; the perpendicular signal in the remaining two quadrants (upper right and lower left) is due to with Hi-C reads mapping equally well to both haplotypes, resulting in spurious interactions. Green lines here are chromosome boundaries.

Urtica genome

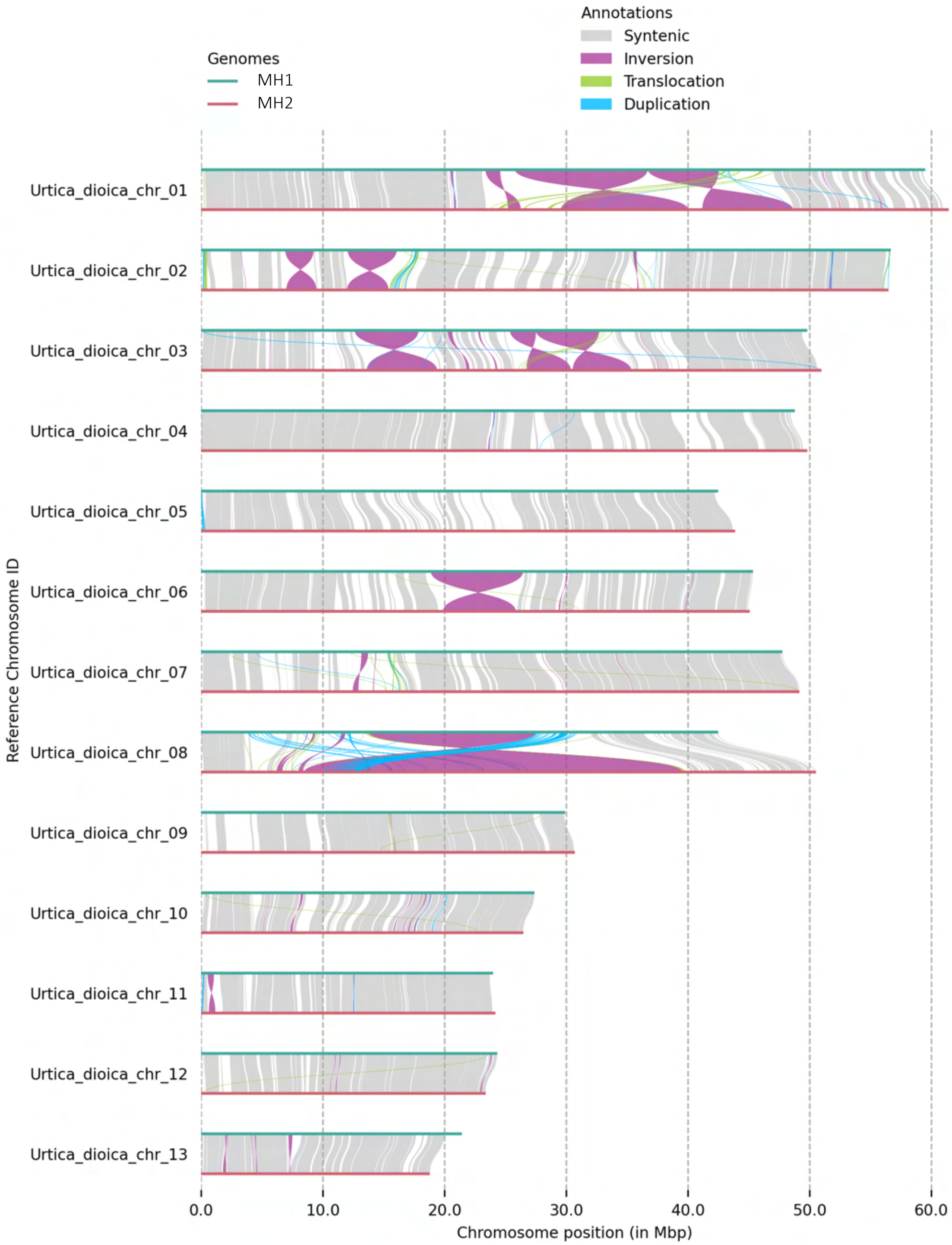


**Supplementary Fig. 2**: **Structural comparison between MH1 and MH2.** Results of comparisons performed with SyRI were visualized using Plotsr.

Urtica genome

24

| Female H1 | Male H1 |
| --- | --- |

Chr1

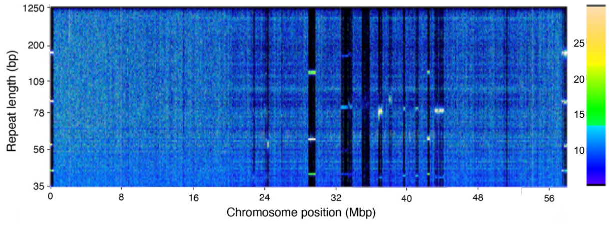


Chr1

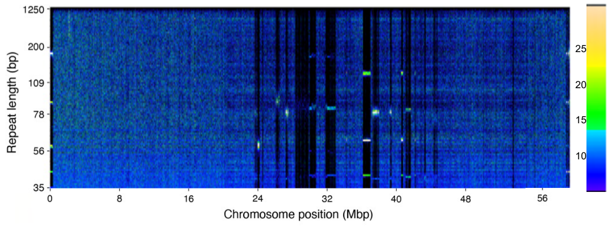


Chr2

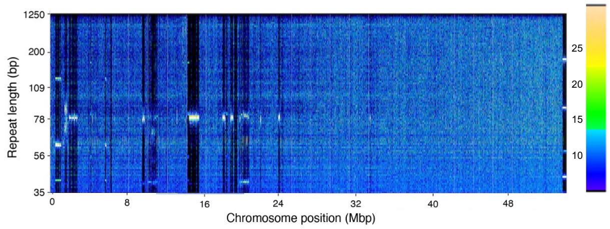


Chr2

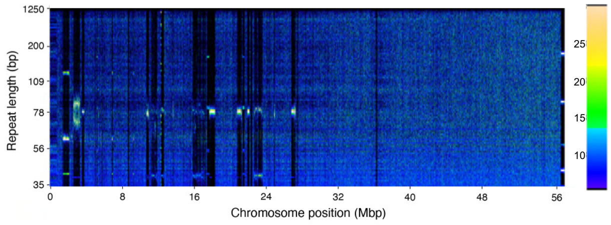


Chr3

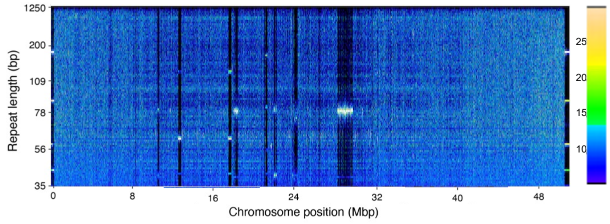


Chr3

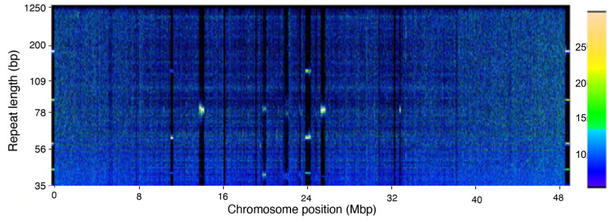


Chr4

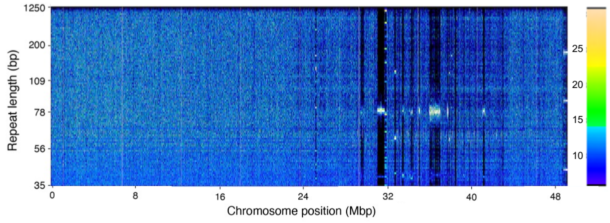


Chr4

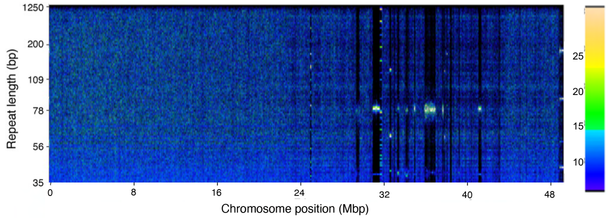


Chr5

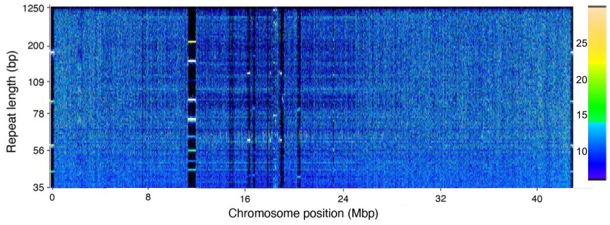


Chr5

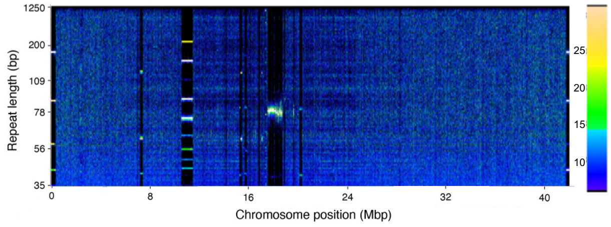

Urtica genome

Chr6

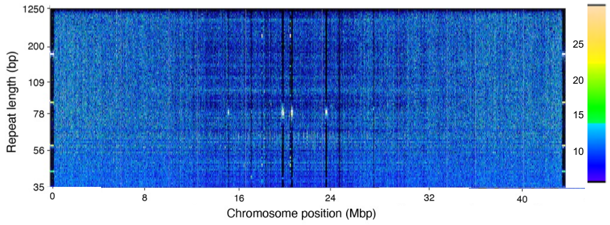


Chr6

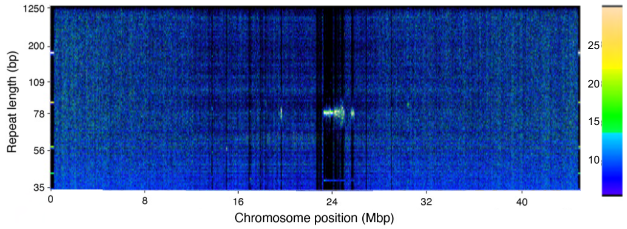


Chr7

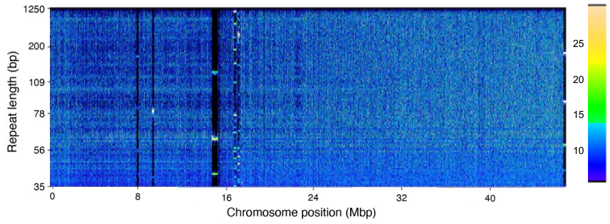


Chr7

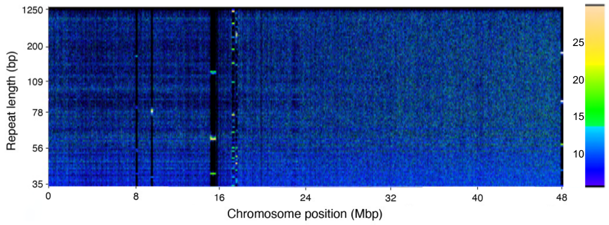


Chr8

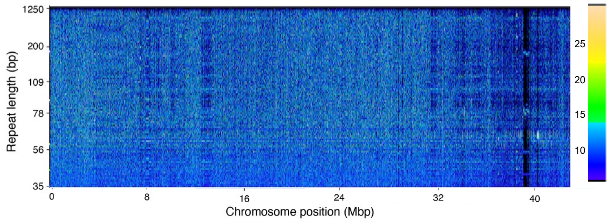


Chr8

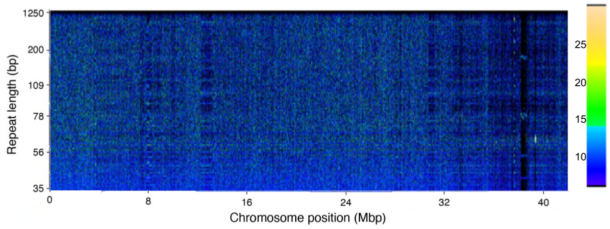


Chr9

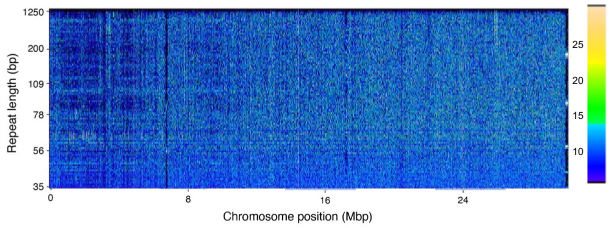


Chr9

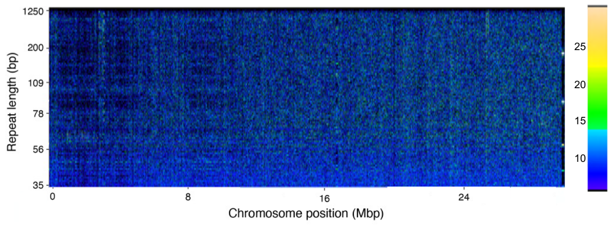


Chr10

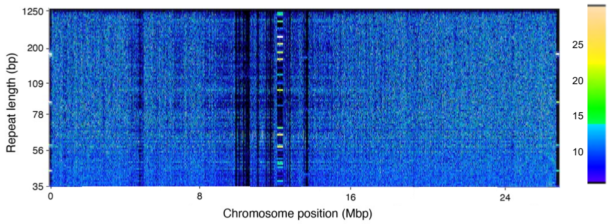


Chr10

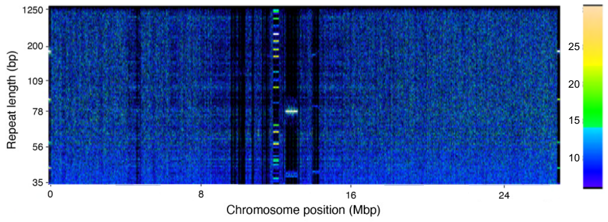

Urtica genome

### Chr11

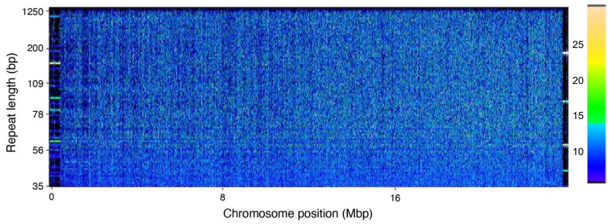


### Chr11

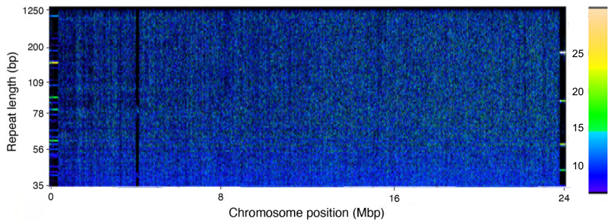


### Chr12

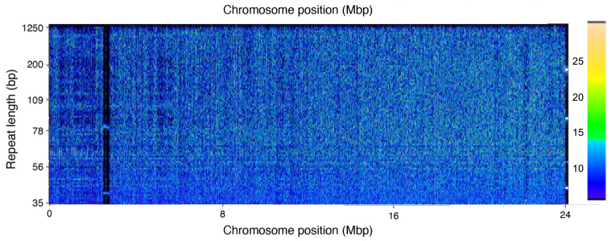


### Chr12

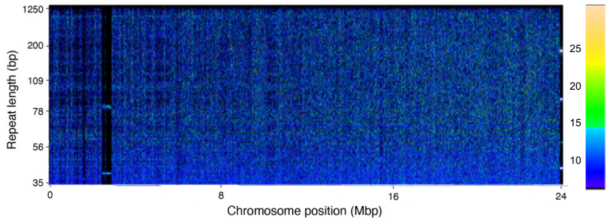


### Chr13

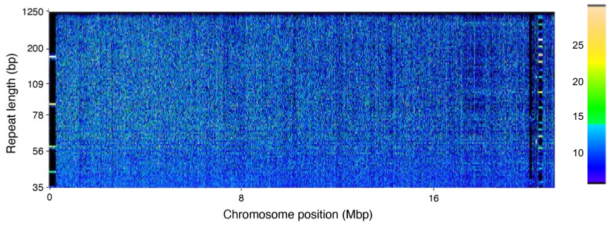


### Chr13

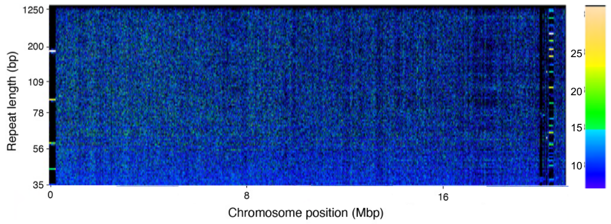


### Female H2

### Male H2

### Chr1

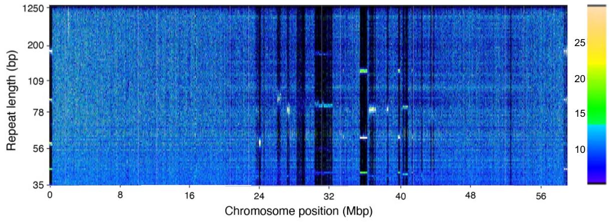


### Chr1

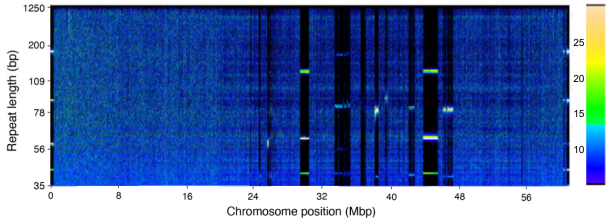


### Chr2

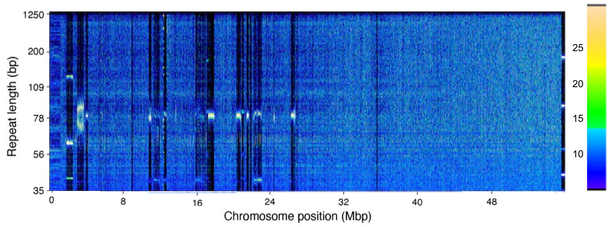


### Chr2

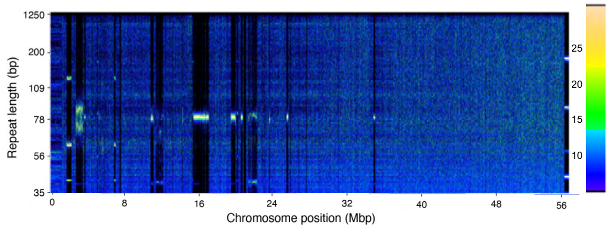

Urtica genome

Chr3

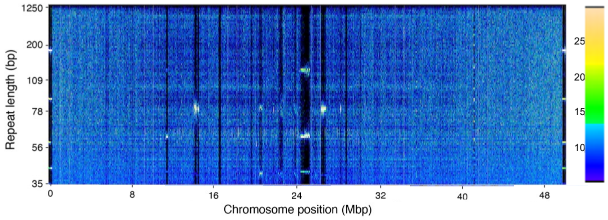


Chr3

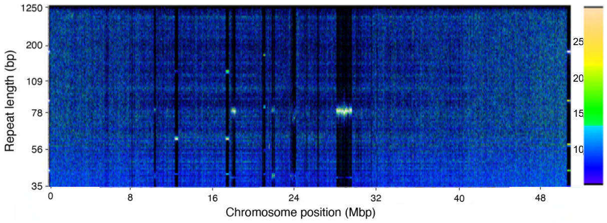


Chr4

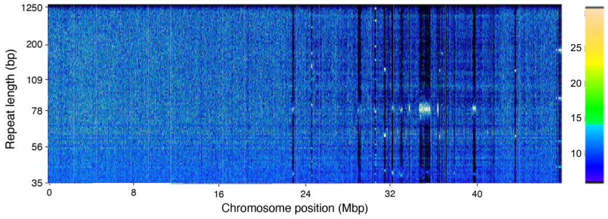


Chr4

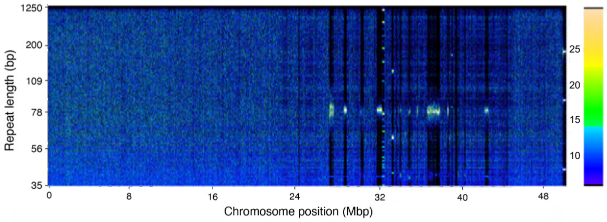


Chr5

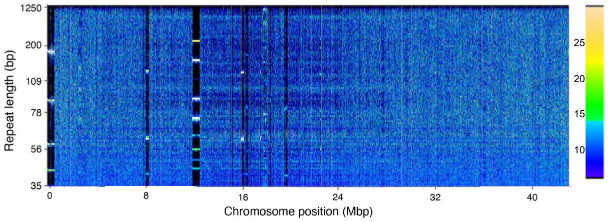


Chr5

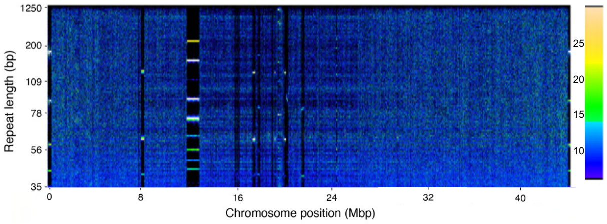


Chr6

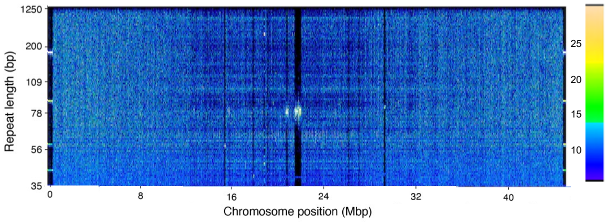


Chr6

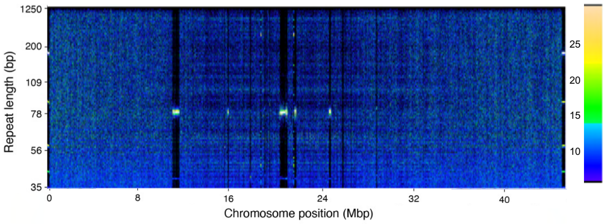


Chr7

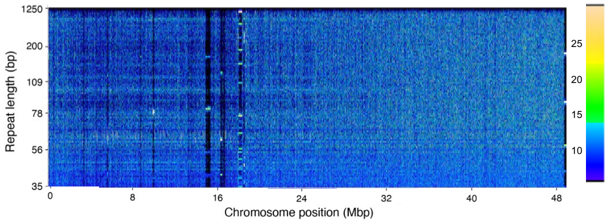


Chr7

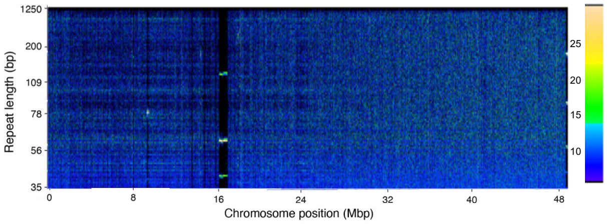

Urtica genome

Chr8

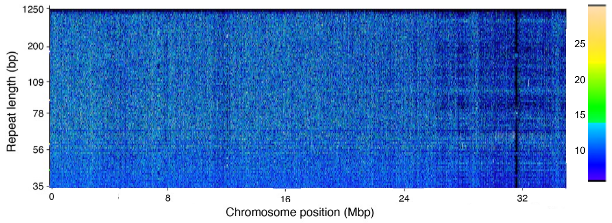


Chr8

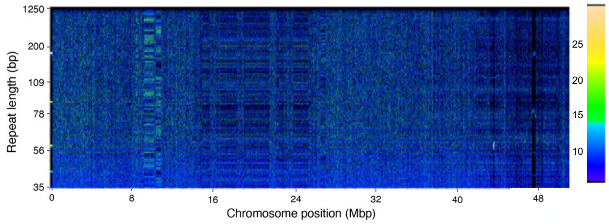


Chr9

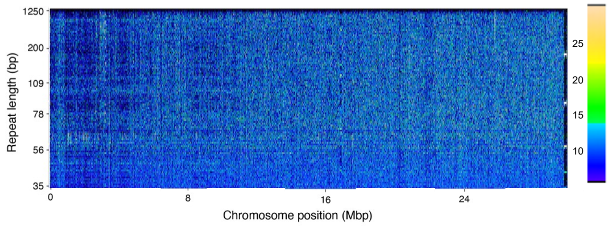


Chr9

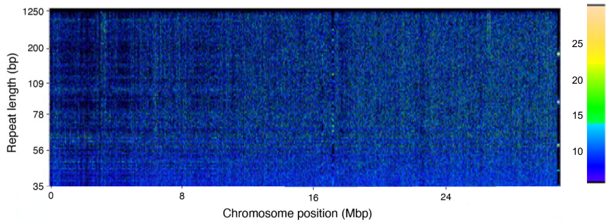


Chr10

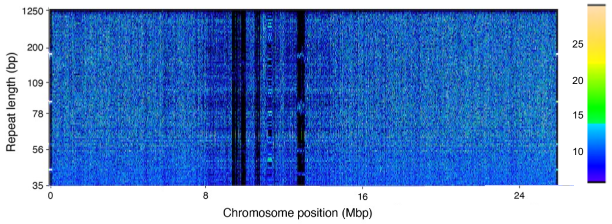


Chr10

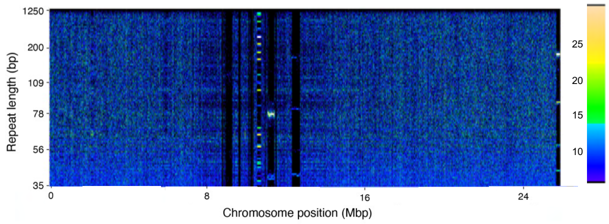


Chr11

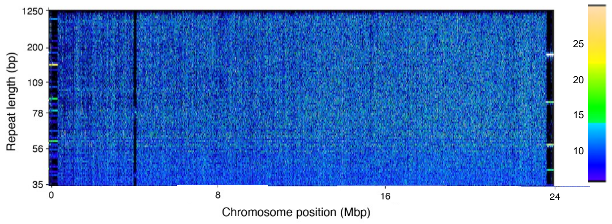


Chr11

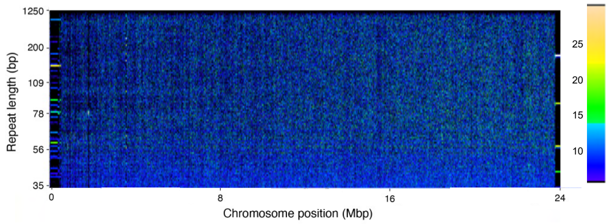


Chr12

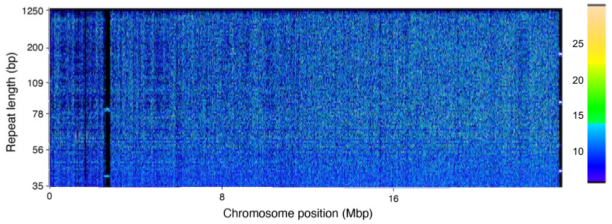


Chr12

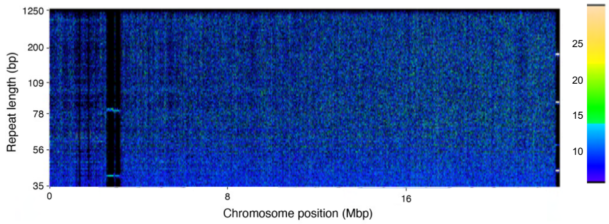

Chr13

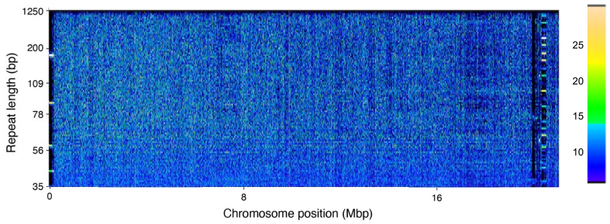


Chr13

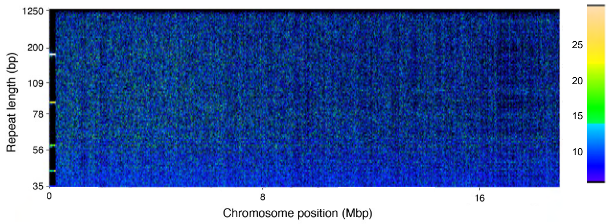


**Supplementary Fig. 3:** **Genome-wide patterns of tandem repeats.** Fourier transformed repeat spectra for each chromosome from the female (left) and male (right) assemblies obtained with RepeatOBserver. The x-axis represents the position on the chromosome, and the y-axis represents repeat length. Colour intensity corresponds to the number of times a specific repeat is found in a 5 kbp window. Regions with high proportions of a specific tandem repeat show characteristic banding patterns (bands corresponding to larger repeats sizes are harmonics of the smaller base repeat).

Chr1

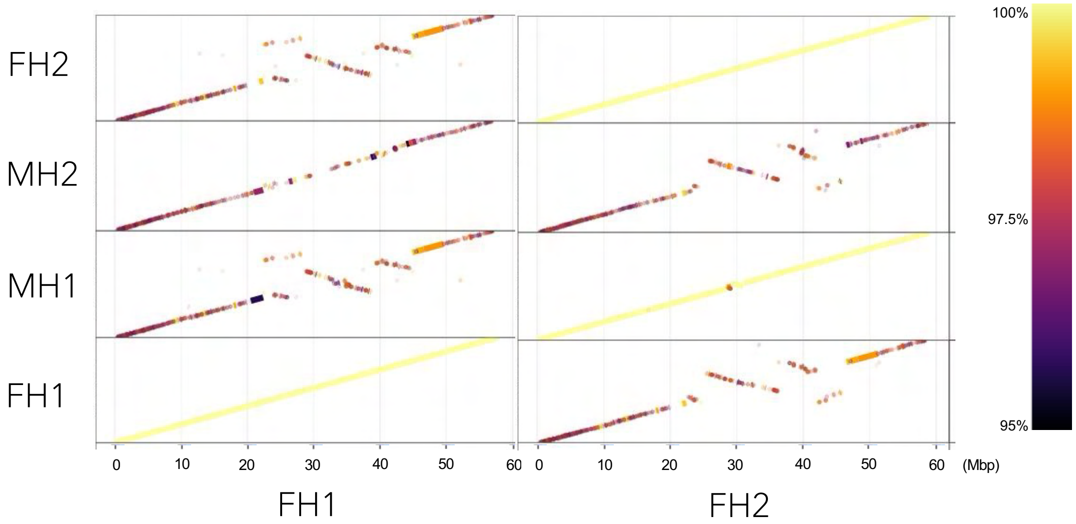


Chr2

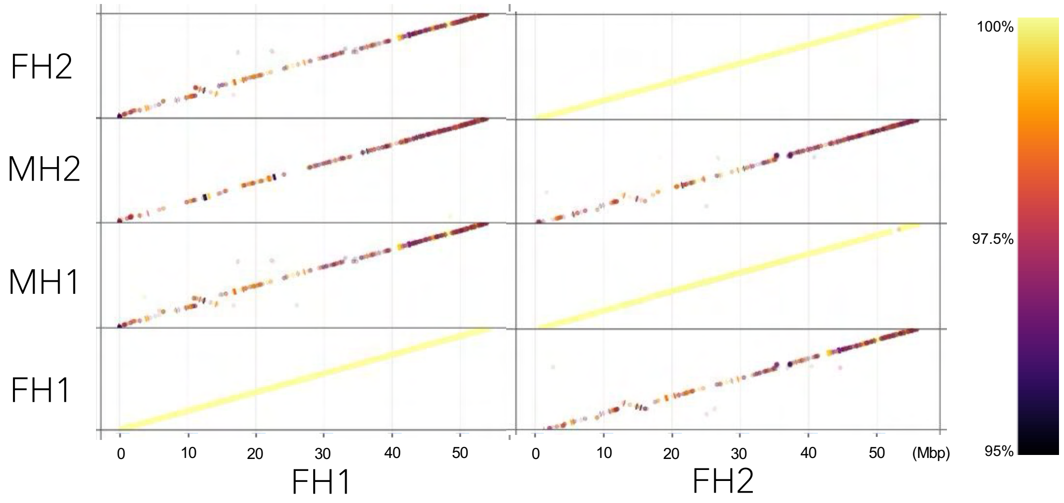


Chr3

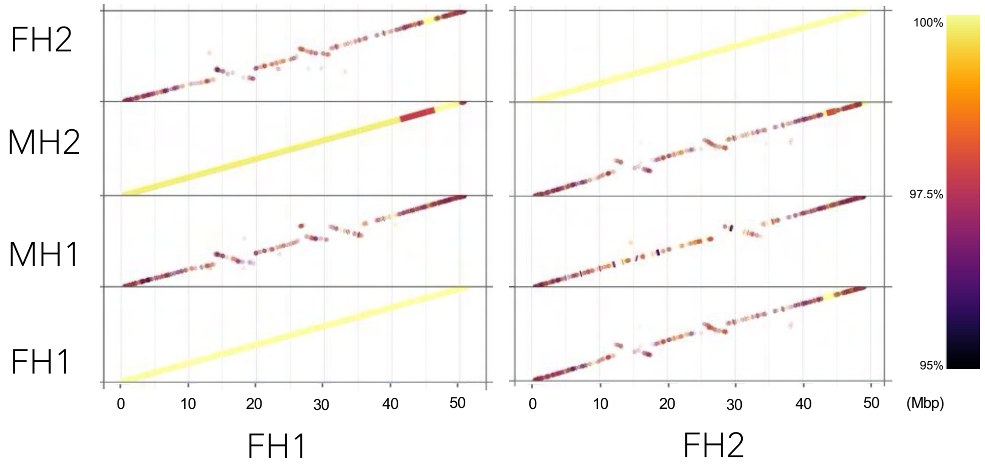

Urtica genome

Chr4

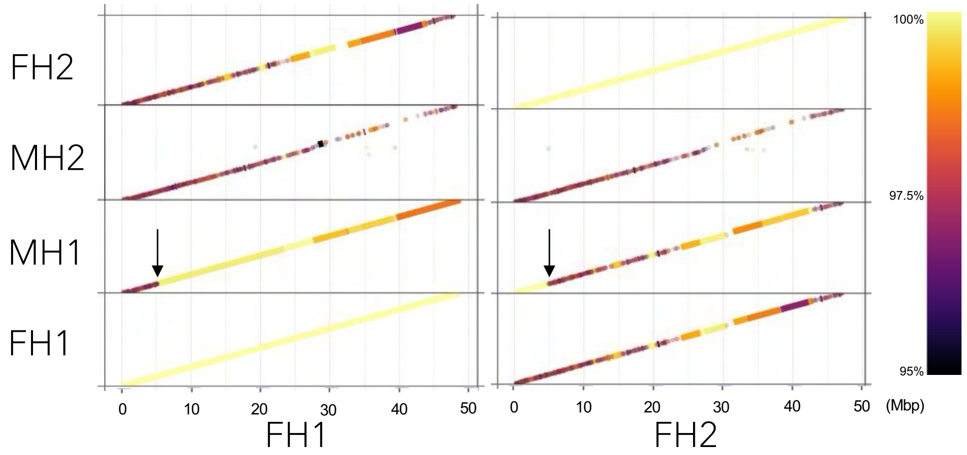


Chr5

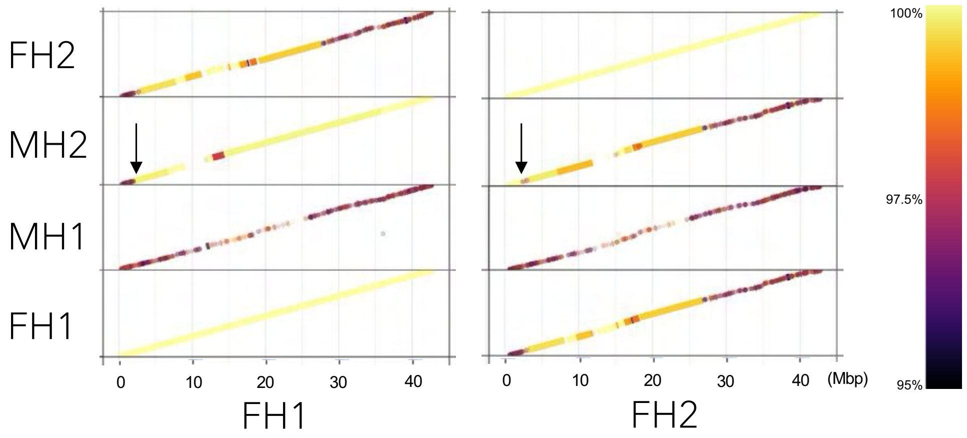


Chr6

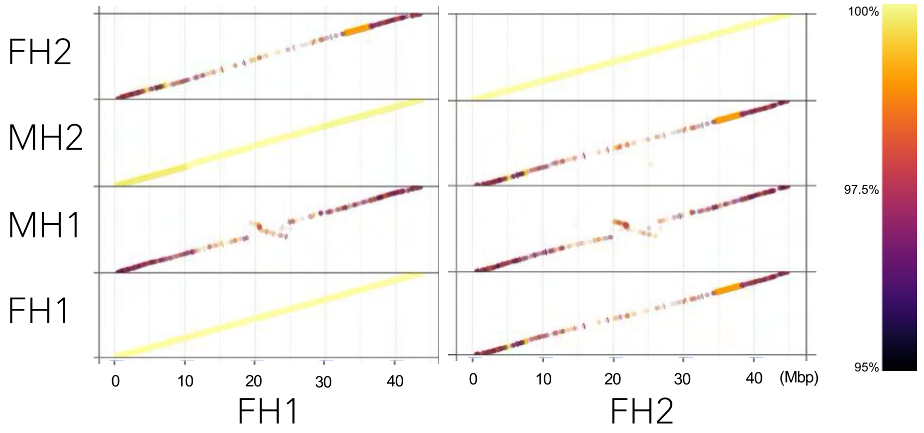

Urtica genome

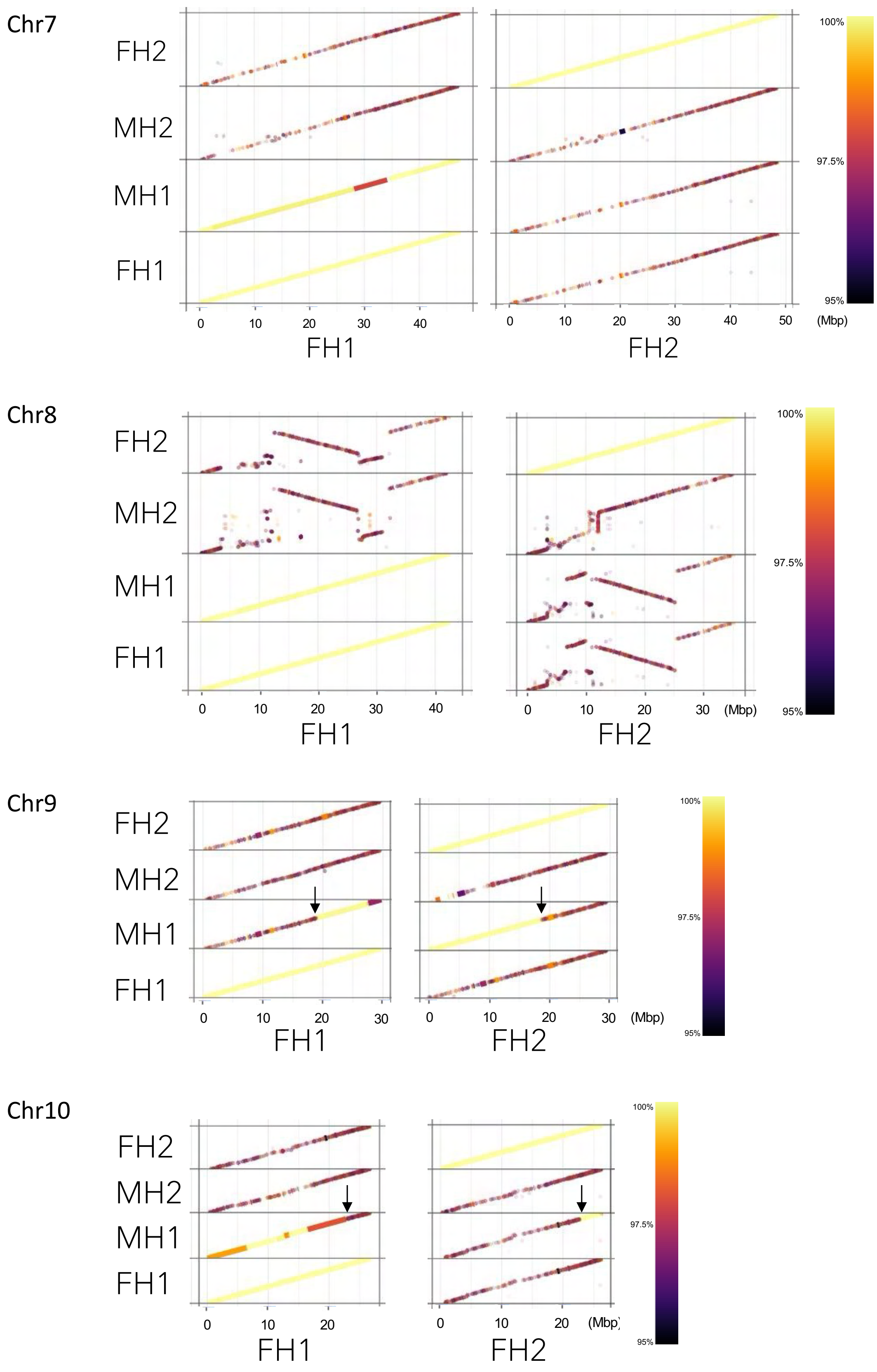

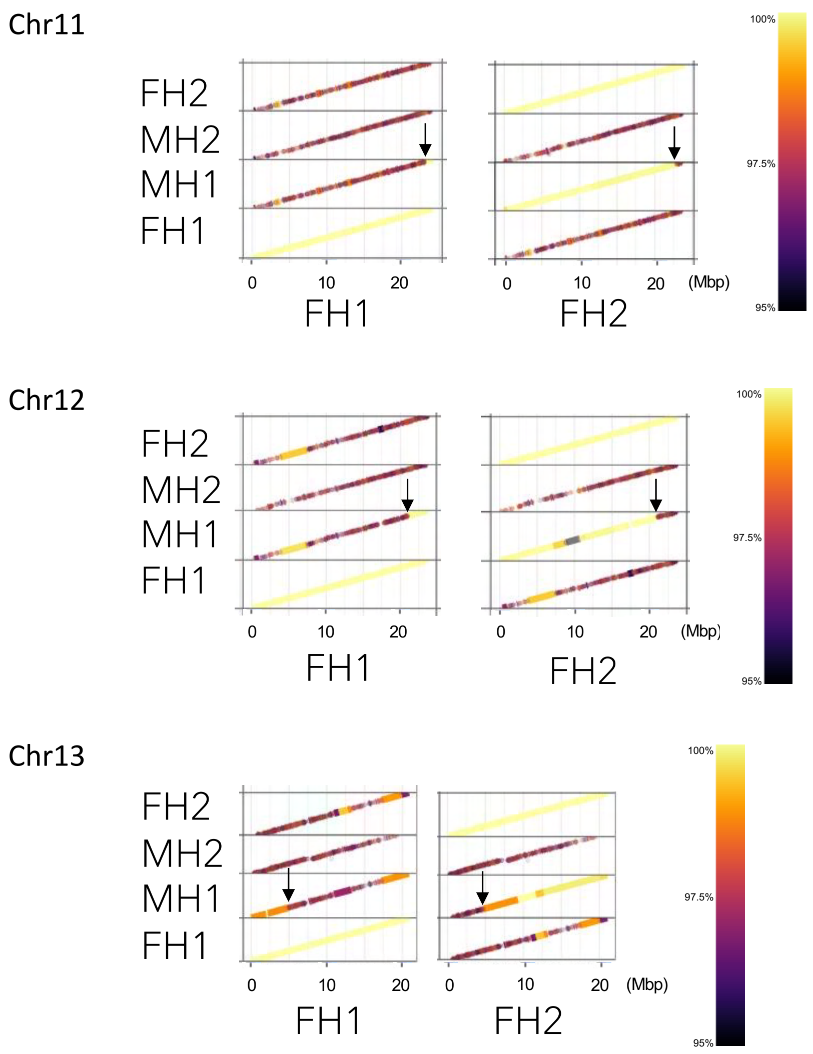


**Supplementary Fig. 4: Maternal and paternal chromosome assignment.** Haplotype similarity and recombination landscapes were obtained by comparing haplotypes from the male and female stinging nettle assemblies. Four haplotypes (FH1 = female haplotype 1, FH2 = female haplotype 2, MH1 = male haplotype 1, MH2 = male haplotype 2) were separately aligned to FH1 and FH2 using minimap2. Our assemblies are haplotype-resolved but not phased. The colours are by nucleotide match in percent identity in gradient scale, and only alignments >95% are shown as a dot/line. The paler the colour, the higher the percent identity of alignment. Yellow = 100% (highest), dark purple = 95% (lowest). Black arrows point at the observed recombination breakpoint based on the switch in the alignment percent pattern between the FH1 and FH2 chromosome inheritance pattern.

**Supplementary Tables**

(Provided on the separate Excel spreadsheet)

**Supplementary Table 1: Genome assembly statistics at different stages of manual curation.**

**Supplementary Table 2: Identification of genes encoding putative pain peptides.**

**Supplementary Table 3: Gene annotation results.**

**Supplementary Table 4: Transposable element annotation results.**

**Supplementary Table 5: Putative centromere positions.**

**Supplementary Table 6: Maternal and paternal chromosome assignment to the male genome assembly based on alignment to the female genome assemblies.**
