## Supplementary Table for "A haplotype-phased male genome sequence of the stinging nettle, *Urtica dioica* ssp. *dioica*"

**Supplementary Table 1: Genome assembly statistics at different stages of manual curation.** The initial genome assembly obtained from Hifiasm went through five stages of curation. (1) Scaffolding with YaHS using Hi-C reads mapped to the MH1 and MH2 assembly individually, with a mapping quality filter >0. (2) Manual correction of obvious misassemblies within MH1 and MH2 using Juicebox. (3) Generation of a haplotype-aware MH1+MH2 Hi-C map, in addition to the individual MH1 and MH2 Hi-C maps, to identify contigs that were assigned to the wrong haplotype (switch errors). To do this, the 3D-DNA pipeline was used, allowing Hi-C reads with mapping quality 0. (4) Removal of potential duplicates and haplotigs, as well as small repetitive sequences that could not be assigned to chromosome scaffolds, with purge_dups with manual thresholds. (5) Final manual check using Juicer, YaHS, and Juicebox. **Sc = scaffold, Cg = contig**

| **Male Haplotype 1** | **Initial draft Hifiasm assembly** | **After scaffolding with YaHS** | **After curation Round 1** | **After curation Round 2** | **After purge_dups** | **After curation Round 3 (final)** |
|---|---|---|---|---|---|---|
| Total length (bp) | 637,325,628 | 637,327,928 | 637,328,928 | 637,419,663 | 576,922,511 | 576,923,711 |
| Scaffold N50 (bp) | NA | 42,714,166 | 42,714,166 | 42,632,247 | 45,393,077 | 45,336,177 |
| Contig N50 (bp) | 29,574,338 | 21,398,079 | 21,398,079 | 13,633,051 | 21,225,977 | 21,136,077 |
| Nr. of contigs/scaffolds | Cg: 1,523 | Sc: 1,514, Cg: 1,537 | Sc: 1,514, Cg: 1,547 | Sc: 1,499, Cg: 1,614 | Sc: 831, Cg: 904 | Sc: 833, Ctg: 918 |
| % main genome in scaffolds >50 kbp | NA | 98.61 | 98.59 | 98.48 | 98.77 | 98.80 |
| Nr. of chromosomes | NA | NA | 13 | 13 | 13 | 13 |
| % genome in chromosome | NA | NA | 81.59 | 81.58 | 90.11 | 90.11 |
| BUSCO (C%) | 93.3 | 93.3 | 93.2 | 93.1 | 92.9 | 93.1 |
| BUSCO (S%) | 89.1 | 89.1 | 89.0 | 88.9 | 90.5 | 90.7 |
| BUSCO (D%) | 4.2 | 4.2 | 4.2 | 4.2 | 2.5 | 2.5 |

| **Male Haplotype 2** | **Initial draft Hifiasm assembly** | **After scaffolding with YaHS** | **After curation Round 1** | **After curation Round 2** | **After purge_dups** | **After curation Round 3 (final)** |
|---|---|---|---|---|---|---|
| Total length (bp) | 556,346,222 | 556,348,122 | 556,354,822 | 556,345,087 | 536,119,383 | 536,126,483 |
| Scaffold N50 (bp) | NA | 45,118,013 | 49,168,886 | 49,119,386 | 49,701,745 | 49,802,645 |
| Contig N50 (bp) | 30,567,971 | 26,435,082 | 15,526,000 | 10,987,956 | 11,166,129 | 11,166,129 |
| Nr. of contigs/scaffolds | Cg: 232 | Sc: 219, Cg: 238 | Sc: 210, Cg: 296 | Sc: 251, Cg: 417 | Sc: 86, Cg: 238 | Sc: 79, Cg: 302 |
| % main genome in scaffolds >50 kbp | NA | 99.92 | 99.84 | 99.54 | 99.65 | 99.80 |
| Nr. of chromosomes | NA | NA | 13 | 13 | 13 | 13 |
| % genome in chromosome | NA | NA | 96.00 | 93.65 | 98.83 | 99.02 |
| BUSCO (C%) | 93.1 | 93.1 | 92.9 | 93.0 | 93.0 | 92.9 |
| BUSCO (S%) | 90 | 90 | 90 | 90 | 90.0 | 89.90 |
| BUSCO (D%) | 3.1 | 3.1 | 3.1 | 3.1 | 3.0 | 3 |

**Supplementary Table 2: Identification of genes encoding putative pain peptides.** The transcript ID from the gene annotation of the stinging nettle male assembly is reported when the region showing homology with pain peptide sequences falls within an annotated gene.

| Name of peptide | Species in which it was originally described | Nt. length | Amino acid length | Source | *U. dioica* male genome position | | | Nr of nt, matches | Nr. of nt in alignment (excl. introns) | Nr, of amino acid matched | Annotated mRNA ID |
|---|---|---|---|---|---|---|---|---|---|---|---|
| | | | | | Chr. | Start | End | | | | |
| **Urthionin A (Δ-Uf1a)** | *Urtica ferox* | 126 | 42 | (Xie et al. 2022) | 09_H1 | 19643373 | 19643499 | 108 | 126 | 36 | NA |
| | | | | | 09_H1 | 19634432 | 19634558 | 108 | 126 | 36 | g13729 |
| | | | | | 09_H2 | 20430702 | 20430828 | 108 | 126 | 36 | g13891 |
| | | | | | 09_H2 | 20421759 | 20421885 | 108 | 126 | 36 | g13890 |
| **Urticatoxin (β/δ-Uf2a)** | *Urtica ferox* | 189 | 63 | (Xie et al. 2022) | 06_H1 | 4513450 | 4513636 | 90 | 198 | 30 | NA |
| | | | | | 06_H2 | 4032458 | 4032644 | 90 | 198 | 30 | NA |
| **Urticatoxin (β/δ-Uf2b)** | *Urtica ferox* | 189 | 63 | (Xie et al. 2022) | 06_H1 | 4513450 | 4513636 | 93 | 198 | 31 | NA |
| | | | | | 06_H2 | 4032458 | 4032644 | 93 | 198 | 31 | NA |
| **Urticatoxin (β/δ-De2a)** | *Dendrocnide excelsa* | 180 | 60 | (Xie et al. 2022) | NA | NA | NA | | | | |
| **Urticatoxin (β/δ-Dm2a)** | *Dendrocnide moroides* | 183 | 61 | (Xie et al. 2022) | 09_H1 | 28134026 | 28134092 | 48 | 66 | 16 | NA |
| | | | | | 09_H1 | 27475565 | 27475631 | 48 | 66 | 16 | NA |
| | | | | | 09_H2 | 28908084 | 28908150 | 48 | 66 | 16 | NA |
| | | | | | 09_H2 | 28377232 | 28377298 | 48 | 66 | 16 | NA |
| **Excelsatoxin A** | *Dendrocnide excelsa* | 105 | 35 | (Gilding et al. 2020) | NA | NA | NA | | | | |
| **Moroidotoxin A** | *Dendrocnide moroides* | 105 | 35 | (Gilding et al. 2020) | NA | NA | NA | | | | |

**Supplementary Table 3. Gene annotation results.** All annotations were done using BRAKER3 v3.0.8 on the softmasked genome, using the mode with RNAseq alignment files and the general plant protein database (Viridiplantae).

| Male Haplotype 1 | BRAKER3 | % of genome |
|---|---|---|
| Total # unique genes annotated | 20,237 | |
| Total # of amino acids/CDS (includes isoforms) | 22,384 | |
| BUSCO (C%) | 90.4 | |
| BUSCO (S%) | 79.7 | |
| BUSCO (D%) | 10.7 | |
| Average length of gene (bp) | 2,553 | |
| Total length of genic region (bp) | 51,673,883 | 8.96 |
| Average length of CDS (bp) | 236 | |
| Total length of coding region (bp) | 28,187,705 | 4.89 |

| Male Haplotype 2 | BRAKER3 | % genome |
|---|---|---|
| total # unique genes annotated | 20,431 | |
| total # of genes identified (includes isoforms) | 22,607 | |
| BUSCO (C%) | 90.5 | |
| BUSCO (S%) | 79.4 | |
| BUSCO (D%) | 11.2 | |
| Average length of gene (bp) | 2,521 | |
| Total length of genic region (bp) | 51,514,561 | 9.61 |
| Average length of CDS (bp) | 236 | |
| Total length of coding region (bp) | 28,267,410 | 5.27 |

**Supplementary Table 4. Transposable elements annotation statistics.** TEs were annotated independently for MH1 and MH2. Note that the genome length here is considered excluding Ns present in the assembly, as per EDTA's default behaviour. A table showing counts of TEs for windows that exceeded the axis maximum in Figure 1c is reported at the bottom.

| **Male Haplotype 1** | **Whole genome (576,893,811 bp total, not counting Ns)** | | | |
|---|---|---|---|---|
| **Class** | **Count** | **bp masked** | **% masked** | **% masked per category** |
| **LINE** | -- | -- | -- | 1.91% |
| I | 971 | 5,929,226 | 1.03% | |
| L1 | 4,127 | 3,838,457 | 0.67% | |
| RTE | 4,138 | 1,218,154 | 0.21% | |
| **LTR** | -- | -- | -- | 47.67% |
| Copia | 114,205 | 89,281,885 | 15.48% | |
| Gypsy | 60,152 | 52,138,084 | 9.04% | |
| unknown | 324,134 | 133,571,214 | 23.15% | |
| **SINE** | -- | -- | -- | 2.73% |
| tRNA | 5,071 | 15,758,139 | 2.73% | |
| **TIR** | -- | -- | -- | 8.70% |
| CACTA | 35,732 | 20,566,330 | 3.57% | |
| Mutator | 65,303 | 18,438,711 | 3.20% | |
| PIF_Harbinger | 8,722 | 3,468,877 | 0.60% | |
| Tc1_Mariner | 827 | 470,983 | 0.08% | |
| hAT | 16,812 | 7,205,989 | 1.25% | |
| **nonTIR** | -- | -- | -- | 2.24% |
| helitron | 41,795 | 12,905,929 | 2.24% | |
| **rDNA** | -- | -- | -- | 0.67% |
| 45S | 3,772 | 3,838,890 | 0.67% | |
| **Repeat fragment** | 39,056 | 22,037,865 | 3.82% | 3.82% |
| **Total interspersed** | **724,918** | **390,681,099** | **67.72%** | |

| **Male Haplotype 2** | **Whole genome (536,073,483 bp total, not counting Ns)** | | | |
|---|---|---|---|---|
| **Class** | **Count** | **bp masked** | **% masked** | **% masked per category** |
| **LINE** | -- | -- | -- | 1.06% |
| I | 33 | 10485 | 0.00% | |
| L1 | 7806 | 7367225 | 1.37% | |
| RTE | 9,927 | 5,684,765 | 1.06% | |
| **LTR** | -- | -- | -- | 46.04% |
| Copia | 58,277 | 58,389,105 | 10.89% | |
| Gypsy | 66,193 | 53,833,182 | 10.04% | |
| unknown | 190,827 | 134,595,808 | 25.11% | |
| **TIR** | -- | -- | -- | 15.08% |
| CACTA | 46,745 | 30,951,459 | 5.77% | |
| Mutator | 92,170 | 39,762,287 | 7.42% | |
| PIF_Harbinger | 7,494 | 2,895,718 | 0.54% | |
| Tc1_Mariner | 295 | 248,758 | 0.05% | |
| hAT | 16,521 | 6,995,186 | 1.30% | |
| **low_complexity** | -- | -- | -- | 0.01% |
| | 214 | 78,424 | 0.01% | |
| **nonTIR** | -- | -- | -- | 2.26% |
| helitron | 37,541 | 12,119,760 | 2.26% | |
| **rDNA** | -- | -- | -- | 0.21% |
| 45S | 3,005 | 1,115,289 | 0.21% | |
| **Repeat fragment** | 34,976 | 9,178,016 | 1.71% | 1.71% |
| **Total interspersed** | **572,024** | **363,225,467** | **67.76%** | |

| **Chromosome** | **Window start** | **Window end** | **TE count** |
|---|---|---|---|
| Urtica_dioica_H1_chr_01 | 36000000 | 36499999 | 7799 |
| Urtica_dioica_H1_chr_01 | 36500000 | 36999999 | 4009 |
| Urtica_dioica_H1_chr_01 | 37000000 | 37499999 | 5156 |
| Urtica_dioica_H1_chr_10 | 12500000 | 12999999 | 4405 |
| Urtica_dioica_H1_chr_10 | 13000000 | 13499999 | 5238 |
| Urtica_dioica_H1_chr_02 | 1500000 | 1999999 | 6674 |
| Urtica_dioica_H1_chr_02 | 2500000 | 2999999 | 4362 |
| Urtica_dioica_H1_chr_02 | 16000000 | 16499999 | 4483 |
| Urtica_dioica_H1_chr_02 | 17500000 | 17999999 | 5581 |
| Urtica_dioica_H1_chr_02 | 20500000 | 20999999 | 4215 |
| Urtica_dioica_H1_chr_02 | 22500000 | 22999999 | 4494 |
| Urtica_dioica_H1_chr_02 | 23000000 | 23499999 | 4629 |
| Urtica_dioica_H1_chr_02 | 26500000 | 26999999 | 4460 |
| Urtica_dioica_H1_chr_03 | 24500000 | 24999999 | 6244 |
| Urtica_dioica_H1_chr_03 | 26000000 | 26499999 | 4035 |
| Urtica_dioica_H1_chr_04 | 31000000 | 31499999 | 4757 |
| Urtica_dioica_H1_chr_04 | 35500000 | 35999999 | 4363 |
| Urtica_dioica_H1_chr_04 | 36000000 | 36499999 | 5316 |
| Urtica_dioica_H1_chr_04 | 40500000 | 40999999 | 4137 |
| Urtica_dioica_H1_chr_05 | 18000000 | 18499999 | 5703 |
| Urtica_dioica_H1_chr_05 | 18500000 | 18999999 | 5602 |
| Urtica_dioica_H1_chr_06 | 23500000 | 23999999 | 6104 |
| Urtica_dioica_H1_chr_06 | 24000000 | 24499999 | 5408 |
| Urtica_dioica_H1_chr_06 | 24500000 | 24999999 | 4699 |
| Urtica_dioica_H1_chr_07 | 15000000 | 15499999 | 7835 |

**Supplementary Table 5. Coordinates of putative centromeric regions.** Centromeric regions as shown in Figure 1e (black bands) are based on the standard deviation cut-off of the repeats Shannon diversity index within the chromosome (Shannon_bin_size=500, Shannon_SD=2) calculated by RepeatOBserver. Compare to Supplementary Figure 3, which shows the tandem repeat pattern in the Fourier transform spectra, and directly visualizes the patterns of tandem repeats across the chromosomes.

| Male Haplotype 1 | | |
|---|---|---|
| Chromosome | Start | End |
| 1 | 27975000 | 30160000 |
| 1 | 35870000 | 37525000 |
| 2 | 1820000 | 3185000 |
| 2 | 16590000 | 18395000 |
| 2 | 21040000 | 21215000 |
| 2 | 21225000 | 22215000 |
| 3 | 15420000 | 15440000 |
| 3 | 23665000 | 23665000 |
| 3 | 24675000 | 27135000 |
| 3 | 48545000 | 48565000 |
| 4 | 29950000 | 32295000 |
| 4 | 35280000 | 35960000 |
| 4 | 36070000 | 37400000 |
| 5 | 17125000 | 19515000 |
| 6 | 22800000 | 25595000 |
| 7 | 8425000 | 9370000 |
| 7 | 14170000 | 16680000 |
| 7 | 46500000 | 46515000 |
| 8 | 37680000 | 40340000 |
| 9 | 1250000 | 1255000 |
| 9 | 1940000 | 3385000 |
| 9 | 3650000 | 3655000 |
| 9 | 3695000 | 3885000 |
| 9 | 28510000 | 28655000 |
| 10 | 11015000 | 11045000 |
| 10 | 11055000 | 11060000 |
| 10 | 11820000 | 13990000 |
| 11 | 1250000 | 1550000 |
| 11 | 22525000 | 22735000 |
| 12 | 1550000 | 3815000 |
| 13 | 1250000 | 1450000 |
| 13 | 19405000 | 20180000 |

| Male Haplotype 2 | | |
|---|---|---|
| Chromosome | Start | End |
| 1 | 34195000 | 34970000 |
| 1 | 43785000 | 46495000 |
| 2 | 2230000 | 3195000 |
| 2 | 15185000 | 17420000 |
| 2 | 20945000 | 20955000 |
| 3 | 27560000 | 27560000 |
| 3 | 27580000 | 30140000 |
| 4 | 31025000 | 31370000 |
| 4 | 31890000 | 33005000 |
| 4 | 36255000 | 38570000 |
| 5 | 11060000 | 13355000 |
| 6 | 10285000 | 12320000 |
| 6 | 19590000 | 21855000 |
| 7 | 15190000 | 17795000 |
| 8 | 1250000 | 1330000 |
| 8 | 41945000 | 44380000 |
| 8 | 45905000 | 48350000 |
| 9 | 1250000 | 1255000 |
| 9 | 1850000 | 3475000 |
| 9 | 3485000 | 3485000 |
| 9 | 29320000 | 29465000 |
| 10 | 10080000 | 11065000 |
| 10 | 11380000 | 12615000 |
| 11 | 1250000 | 1690000 |
| 11 | 22755000 | 22945000 |
| 12 | 1795000 | 3985000 |
| 13 | 1250000 | 1455000 |

**Supplementary Table 6. Maternal and paternal chromosome assignment to the male genome assembly based on alignment to the female genome assembly** (Hirabayashi et al. 2025). Maternally-inherited chromosomes were identified based on visually inspecting the pairwise alignment plot coloured by the percent nucleotide identity (Figure 2, Supplementary Figure 4), where FH1 and FH2 were used as a reference genome. More complete alignment between haplotype pairs was considered as evidence of maternal inheritance. To calculate the proportion of a male haplotype (MH1/MH2) in perfect alignment with the female haplotypes (FH1/FH2), we used >99.7% alignment blocks and calculated the (total length of alignment / total chromosome length x 100%) for each chromosome pair.

| | Proportion of MH1 aligning with | | Proportion of MH2 aligning with | | Male haplotypes | |
|---|---|---|---|---|---|---|
| **Chromosome #** | **FH1** | **FH2** | **FH1** | **FH2** | **Maternal haplotype** | **Paternal haplotype** |
| 1 | 1.2% | 98.4% | 0.6% | 0.2% | MH1 | MH2 |
| 2 | 2.6% | 97.4% | 0.0% | 0.2% | MH1 | MH2 |
| 3 | 0.0% | 0.0% | 87.3% | 4.8% | MH2 | MH1 |
| 4 | 46.4% | 23.1% | 1.6% | 0.3% | MH1 | MH2 |
| 5 | 0.0% | 0.1% | 84.8% | 19.2% | MH2 | MH1 |
| 6 | 0.0% | 0.0% | 99.6% | 2.1% | MH2 | MH1 |
| 7 | 86.2% | 1.3% | 0.1% | 0.0% | MH1 | MH2 |
| 8 | 99.5% | 0.5% | 0.3% | 0.3% | MH1 | MH2 |
| 9 | 29.9% | 63.6% | 0.2% | 0.9% | MH1 | MH2 |
| 10 | 34.1% | 13.6% | 1.0% | 0.0% | MH1 | MH2 |
| 11 | 7.5% | 93.1% | 0.2% | 0.2% | MH1 | MH2 |
| 12 | 11.6% | 72.0% | 0.0% | 0.1% | MH1 | MH2 |
| 13 | 0.8% | 47.8% | 0.0% | 0.0% | MH1 | MH2 |
